# Repeated restraint stress-induced increase in post-surgical somatosensory hypersensitivity and affective responding is mediated by β-adrenergic receptor activation and spinal NLRP3-IL1β signalling in male rats

**DOI:** 10.1101/2025.05.06.652397

**Authors:** Ariadni Bella, Khaled Abdallah, Daniela Rodrigues-Amorim, Alba M. Diego, Charles Decraene, Pierre Hener, Chiara Di Marino, David P. Finn, Ipek Yalcin, Michelle Roche

## Abstract

Pre-surgical stress is a well-recognised risk factor for persistent post-surgical pain, and while the precise underlying neurobiological mechanisms remain unknown, neuro-immune interactions are believed to play a pivotal role. Here, we investigated the effect of repeated restraint stress (RRS) on post-surgical somatosensory hypersensitivity and affective responding in male rats and examined underlying mechanisms mediating these effects. We showed that RRS induced behavioural despair in the forced swim test, reduced body weight gain and elevated faecal corticosterone levels in male Sprague-Dawley rats. Following paw-incision surgery, animals pre-exposed to RRS exhibited exacerbated mechanical and heat hypersensitivity, pain-related aversion, and anxiety-like behaviour compared to non-stress counterparts. RNAseq analysis revealed alterations in glial and neuro-immune pathways in the dorsal horn of the spinal cord in the RRS + paw incision group compared to paw incision alone, data further confirmed by increased microglial activity and inflammatory gene expression (*iba1*, *itgam*, *il-1β* and *nlrp3*). Intrathecal administration of IL-1Ra or MCC950 (an NLRP3 inhibitor) attenuated the RRS-induced increase in pain-related aversion and mechanical hypersensitivity post-surgery. Chronic administration of RU486, a glucocorticoid receptor antagonist, prevented RRS-induced despair-like behaviour but did not alter the effects of RRS on pain-related aversion or mechanical hypersensitivity post-surgery. In contrast, chronic administration of propranolol, a β-adrenergic receptor antagonist and sympathetic nervous system inhibitor, not only prevented the RRS-induced despair-like behaviour but also attenuated exacerbation of mechanical hypersensitivity, pain-related aversion, and anxiety-like behaviour post-surgery. These findings suggest that RRS exacerbates and prolongs post-surgical somatosensory and affective pain responding via β-adrenergic receptor activation and increased spinal microglial NLRP3-IL1β signalling. These data provide further insight into the mechanisms by which chronic stress and mood disorders exacerbate and prolong post-surgical pain.

## 1. INTRODUCTION

Each year, over 300 million people undergo surgery worldwide (Richebe et al., 2018). While acute post-surgical pain is an expected and manageable outcome, 10-50% of patients develop prolonged and persistent post-surgical pain (PPP) (Fuller et al., 2023). This transition from acute to persistent pain presents a significant clinical challenge, often complicated by psychological factors and pre-existing mood disorders, such as depression, which is linked to greater pain intensity and delayed recovery (AbuRuz, 2019; Dunn et al., 2018; Gohari et al., 2022; Poole et al., 2017). Despite growing interest in the link between emotional dysregulation and post-surgical pain, our understanding of the mechanisms underlying these interactions remains limited, highlighting the need for further research to improve patient outcomes.

Animal models, such as the hind paw incision model, have provided critical insights into the mechanisms underlying post-surgical pain. In this model, animals exhibit mechanical and heat hypersensitivity (Brennan et al., 1996), as well as anxiety-like behaviour (Dai et al., 2011; Li et al., 2010), which typically resolve within five to seven days. However, exposure to stress prior to surgery can result in more intense and prolonged post-surgical sensory hypersensitivity. Acute stress in the form of a single prolonged stress (SPS) episode resulted in prolonged mechanical hypersensitivity for up to 35 days post-surgery (Liu et al., 2015; Sun et al., 2019; Wu et al., 2019; Zhang et al., 2019). Similarly, pre-operative REM sleep disturbance (RSD) (Li et al., 2019; Wang et al., 2015; Xue et al., 2018) or repeated immobilization stress for 3 days (Cao et al., 2015) increased mechanical and heat hypersensitivity in male rats for up to 9 days post paw-incision surgery. Chronic stress paradigms such as repeated restraint/immobilisation or repeated defeat stress (RDS) increased the magnitude and duration of post-surgical mechanical hypersensitivity for between 9-24 days (Aizawa et al., 2018; Aizawa et al., 2019; Arora et al., 2018; Li et al., 2014)(Meng et al., 2021). However, the effect of stress on other pain-related outcomes such as heat hypersensitivity, aversive behaviour and affective responding is unknown.

Several mechanisms may underlie the effects of stress on the prolongation of post-surgical pain [for review see (Bella et al., 2022)], including spinal neuro-immune mechanisms. For example, SPS has been found to increase proinflammatory cytokine levels and lead to spinal astrocyte activation for up to 28 days post-surgery (Liu et al., 2015), while inhibition of astrocyte activation was able to prevent the exacerbation of mechanical hypersensitivity post-surgery (Zhang et al., 2019). Furthermore, minocycline (a microglia inhibitor) (Sun et al., 2017), an alpha7 nicotinic acetylcholine receptor (α7 nAChR) antagonist (Sun et al., 2019) or GSK650394 (a serum- and glucocorticoid-regulated kinase 1 (SGK1) inhibitor) (Zhang et al., 2019), attenuated SPS-induced potentiation of mechanical hypersensitivity and spinal neuroinflammation. In relation to chronic stress, increased NLRP3-IL-1β signalling in the basolateral amygdala (BLA) has been shown to mediate the prolongation of post-surgical mechanical hypersensitivity following chronic repeated restraint stress (Meng et al., 2021). However, it is unknown if spinal neuroinflammatory mechanisms may also play a role in the effects of chronic stress on post-surgical pain responding.

Stress is well recognised for its ability to prime microglia, creating a neuro-immune environment more susceptible to heightened responses upon subsequent challenges (de Pablos et al., 2006; Espinosa-Oliva et al., 2011; Frank et al., 2007; Frank et al., 2018; Frank et al., 2020; Johnson et al., 2004; Wohleb et al., 2011). Activation of the two key stress pathways, the hypothalamic–pituitary–adrenal axis (HPA axis) and the sympathetic nervous system (SNS), has been shown to increase microglia activity, leading to increased production of proinflammatory mediators such as IL-1β and TNF-α (Feng et al., 2019; Frank et al., 2012; Munhoz et al., 2006; Wohleb et al., 2011). For example, Feng and colleagues demonstrated that repeated restraint stress for 21 days increases the expression of *iba1*, *il-1β* and *il-18* in hippocampal microglia and induces depressive-like behaviour, effects mediated by activation of the glucocorticoid receptor (GR)-NF-κB-NLRP3 signalling pathway (Feng et al., 2019). In addition, antagonism of the GR has been shown to prevent chronic restraint stress-induced neuroinflammation in the spinal cord (Meyer et al., 2024). Inhibition of SNS activation/β-adrenergic activation has been shown to block repeated social defeat-induced anxiety-like behaviour and enhanced brain neuroinflammation (Wohleb et al., 2011). Thus, glucocorticoid and β-adrenergic receptors play a pivotal role in mediating chronic stress-induced neuroinflammation and may underlie the effects of chronic stress on post-surgical pain responding.

We hypothesised that chronic stress would result in glucocorticoid and/or β-adrenergic receptor activation and spinal microglial priming, resulting in enhancement of NLRP3-IL-1β signalling, prolongation of mechanical and heat hypersensitivity and exacerbation of affective pain responding post surgery. To test this hypothesis, we used repeated restraint stress (RRS) combined with the hind paw incision model of post-surgical pain to develop a model of prolonged post-surgical hypersensitivity and affective responding. A combination of molecular and pharmacological approaches was then used to uncover the mechanisms involved. We found that RRS markedly amplifies both sensory hypersensitivity and affective pain following surgery, effects mediated by β-adrenergic receptor activation and spinal NLRP3-IL-1β signalling.

## 2. METHODS

### 2.1 Animals

A total of 288 adult male Sprague-Dawley rats (170-220 g upon arrival; Janvier, France) were used for the experiments. Male rats were used in this study as pilot data from our group has demonstrated that RRS-induced prolongation of post-surgical pain behaviours were not associated with increases in the spinal expression of *iba1*, *nlrp3* or *il-1β* in female rats, suggesting distinct sexual dimorphic mechanisms. Animals were housed in groups of two or three in type IV cages (55×33×20cm) under a 12:12 h light–dark cycle (lights on at 8am) and maintained at controlled conditions (temperature 22 ± 2 ◦C, humidity 45–55%), with food and water ad libitum. All experiments were carried out during the light phase. The experimental procedures were approved by the Animal Care and Research Ethics Committee, National University of Ireland Galway and carried out under license from the Health Products Regulatory Authority in the Republic of Ireland and approval by the regional ethical committee of Strasbourg (CREMEAS, APAFIS 43704) in accordance with EU Directive 2010/63 and the ARRIVE guidelines.

### 2.2 Experimental design

A graphical representation of the experimental design is provided in Supplementary Figure 1 (S1). For all experiments, animals were assigned randomly to experimental groups.

#### 2.2.1 Establishment of the chronic stress-induced potentiation of post-surgical somatosensory hypersensitivity model

Rats were exposed to RRS or no RRS for 6hr/day 3 day, 2.5h/day for 14 days or 6hr/day for 21 days. Body weight gain, mechanical and heat sensitivity were assessed throughout the RRS procedure, and the forced swim test (FST) was conducted following the last RRS session. Faecal pellets were collected from a separate cohort of animals that underwent the same procedures. The choice of restraint period was based on the literature demonstrating that these protocols elicit stress-induced alterations in behaviour and/or modulate post-surgical hypersensitivity (Imbe et al., 2012; Meng et al., 2021). On the day following the end of RRS, animals underwent paw incision or sham surgery. Mechanical and heat sensitivity were assessed for up to 3 weeks following paw incision or sham surgery.

#### 2.2.2 The effect of RRS on aversive- and anxiety-like behaviour post-surgery

Rats were exposed to RRS (6h/day 21day) or no stress and the day following the end of RRS/no stress, they underwent paw incision or sham surgery followed by a battery of behavioural aversive or affective tests in the following order (days post-surgery): PEAP (day 2), OFT (day 4), EPM (day 6) and LDB (day 10). Mechanical hypersensitivity was assessed throughout the post-surgical period (days 0-13).

#### 2.2.3 Transcriptomic and molecular alterations in the dorsal horn of the spinal cord to understand mechanisms underlying RRS-induced exacerbation of post-surgical pain

Rats underwent RRS (6h/day 21day) or no stress followed by paw incision or sham surgery. Von Frey and Hargreaves testing was conducted on Day 5 post-surgery, immediately after which animals were euthanised or perfused and L4-L6 dorsal horn was carefully removed and stored for subsequent transcriptomic or immunohistochemical analysis.

#### 2.2.4 The role of the spinal NLRP3/IL-1β pathway in RRS-induced exacerbation of post-surgical mechanical hypersensitivity and aversion

Rats underwent RRS (6h/day 21day or no stress) followed by paw incision surgery. Animals received an intrathecal injection of either the NLRP3 antagonist MCC950 (20μg/30μl; R&D Systems, Inc., Minneapolis, USA), IL-1 receptor antagonist (IL-1Ra) (30μg/30μl Anakinra; human recombinant IL-1Ra, Swedish Orphan Biovitrum) or corresponding vehicle (5% DMSO or Saline). The doses were selected based on the literature (Sanchez et al., 2019; Zhang et al., 2022) and on pilot studies demonstrating that these doses are not analgesic in animals subjected to paw incision in the absence of stress. MCC950, IL-1Ra or the corresponding vehicle were administered intrathecally and effects on pain-related aversion (PEAP, day 2) and mechanical hypersensitivity (Von Frey, day 5) post-surgery were examined 15 minutes following administration.

#### 2.2.5 Determining if the effects of RRS on post-surgical sensory and affective pain responding are due to stress-induced glucocorticoid or β-adrenergic receptor activation

Animals received systemic administration of the glucocorticoid receptor antagonist RU486 (10mg/kg i.p. Merck, Ireland), propranolol hydrochloride (10mg/kg i.p; Merck, Ireland) or vehicle (saline) 30min prior to each RRS or non-stress on each day for 21 days. This dosage and timing were based on previous studies (Dong et al., 2017; Peng et al., 2015). Despair-like behaviour was assessed at the end of the RRS/non stress period (FST) after which all animals underwent paw incision surgery. Mechanical hypersensitivity was assessed on days 1,3,5,7, and 10, post-surgical aversion (PEAP) assessed on day 2 and anxiety-like behaviour (OFT) on day 4.

### 2.3 Repeated restraint stress

In order to establish a model of stress-induced persistent post-surgical pain, we employed three different protocols of Repeated Restraint Stress (RRS). In specific, rats were placed in cylindrical plastic restraining tubes for 3 days (6h/day), 14 days (2.5h/day) or 21 days (6h/day). The choice of restraint period was based on pilot studies and the literature demonstrating that these protocols elicit stress-induced alterations in behaviour and/or modulate post-surgical hypersensitivity (Imbe et al., 2012; Meng et al., 2021).

### 2.4 Plantar hind paw incision

The plantar hind paw incision surgery was conducted in accordance with previous published papers (Brennan et al., 1996). Briefly, animals were anaesthetised with isoflurane. The plantar surface of the left hind paw received a 1-cm longitudinal incision through skin and fascia and the plantaris muscle was lifted and incised longitudinally. After haemostasis, the skin was closed with two 5-0 silk mattress sutures (ETHICON, San Lorenzo, Puerto Rico, USA) and rats were placed in a heated clean cage until recovery. The animals subjected to sham surgery underwent isoflurane anaesthesia for an equivalent time (10-15 min).

### 2.5 Mechanical and Heat Sensitivity

#### 2.5.1 Von Frey test

In order to assess mechanical sensitivity, paw withdrawal thresholds (PWT, g) were determined using an electronic von Frey (IITC, CA, U.S.A.). The stimulus was applied to the mid outer part of the incisional site. Two PWTs were taken per paw, which were then averaged for statistical analysis.

#### 2.5.2 Hargreaves’ test

Immediately following Von Frey, heat sensitivity (paw withdrawal latency, PWL, s) was assessed using the Hargreaves’ test (IITC, CA, USA). PWLs were measured three times/paw with a cutoff of 20s and the average of the three recordings was used for statistical analysis.

### 2.6 Affective Responding

#### 2.6.1 Forced swim test

The two-day FST took place 2 hours after the last RRS session, and the following day as previously described (Porsolt et al., 1977). The behavioural response of the rats (immobility) was assessed blindly by a trained observer with the aid of Ethovision 15.0 XT (Noldus Technology, Wageningen, The Netherlands).

#### 2.6.2 Open Field Test

The Open Field Test (OFT) was conducted as previously described (Thorton, 2021). The open field apparatus consisted of a circular arena (diameter 75 cm) with high walls (40 cm in height) and an inner bright central arena (diameter 40 cm). Each rat was placed in the centre of the arena and allowed to freely explore for a duration of 5 minutes. The time spent in the central zone was recorded using Ethovision 15.0 XT software package (Noldus Technology, Wageningen, The Netherlands).

#### 2.6.3 Elevated Plus Maze

The Elevated Plus Maze (EPM) test was conducted as previously described (Thorton, 2021). Each rat was placed in the central platform facing one of the open arms (75 lux) and allowed to explore the maze for 5 minutes. The time spent in the open arms was recorded and analysed using the Ethovision 15.0 XT software package (Noldus Technology, Wageningen, The Netherlands).

#### 2.6.4 Light-Dark Box

Light/Dark Box (LDB) test was conducted as previously described (Becker et al., 2023). The apparatus consisted of two connected compartments (40 cm × 40 cm x 60cm) either brightly lit (1000 lux) or dark (<10 lux). On the test day, rats were placed in the dark compartment at the start of the trial, and their behaviour was recorded in the test arena for 5 minutes using a video camera positioned above the apparatus. Ethovision 15.0 XT software package (Noldus Technology, Wageningen, The Netherlands) was used to analyse the time spent in each compartment.

#### 2.6.5 Emotionality z-scores

To evaluate emotional behaviour across multiple tests, a composite emotionality Z-score was calculated using data from the OFT, EPM and LDB. For each test, raw scores (e.g., time spent in the centre of the arena, time spent in the open arms, time spent in the bright compartment) were converted into Z-scores to standardize the data across measures. The Z-scores were computed as follows: Z=(X−μ) /σ, where X is the individual score from a given test, μ is the mean of the scores within the control group and σ is the standard deviation of the control group (Guilloux et al., 2011). Z-scores from each behavioural test were then averaged to generate an emotionality score for each animal, reflecting the overall level of emotional behaviour.

### 2.7 Place Escape/Avoidance Paradigm

The affective component of pain or pain-related aversion was assessed using the Place Escape/Avoidance Paradigm (PEAP) (LaBuda & Fuchs, 2000). In brief, the PEAP apparatus consisted of a brightly lit (1000 lux) and dark chamber (40cm x 40cm x 60cm each compartment). Rats were placed in the dark compartment and allowed to freely explore both compartments for a total duration of 30 minutes during which time a 15g Von Frey filament was applied to either the ipsilateral (when in the dark compartment) or contralateral (when in the light compartment) paw every 15 s. Time spent (s) in the bright compartment, was recorded and used as an indicator of the aversive response to mechanical stimulation.

### 2.8 Corticosterone level measurement

Faecal samples were collected from individual rats 2-3 hours following the last RRS session in accordance with the literature on the peak faecal corticosterone level post-stress (Rowland & Toth, 2019). Corticosterone levels were quantified using the Enzo Life Sciences Corticosterone ELISA Kit (Enzo Life Sciences, Farmingdale, NY) following the manufacturer’s instructions. Results were expressed as ng/g.

### 2.9 Bulk 3’RNA-seq and analysis

RNA was isolated from dorsal horn of the spinal cord using the Nucleospin® RNA II total isolation kit (Macherey-Nagel, Germany). RNA Integrity Number (RIN) values of the samples used are presented in Supplementary Table S2. The 3’RNA-Seq process used is an in-house adaptation of the method developed by Foley and colleagues (Foley et al., 2019), performed by IntegraGen, Paris, France. In summary, 10 ng of total RNA was fragmented and the 3’ ends of mRNA were captured using an RT primer containing a poly(T) sequence and a Unique Molecular Identifier (UMI). Illumina adapters were incorporated during this step via template switching. The resulting fragments underwent two rounds of PCR amplification to complete the Illumina adapters and add indexing sequences. The read structure is as follows: Read 1: Contains a 26-base UMI followed by the beginning of the poly(T) sequence. Read 2: Starts with a GGG sequence (to be trimmed), followed by the variable-length 3’ insert tag and the poly(A) tail. Sequencing was performed on an Illumina NovaSeq 6000 platform using a 2×100 bp paired-end configuration.

The bulk RNA-seq data analysis was performed using Galaxy (Galaxy, 2024). Preprocessing began using Trimmomatic, where adapter sequences were removed, and reads were trimmed if their quality scores fell below 20. Reads shorter than 35 base pairs were discarded. Quality control was subsequently performed using FastQC, ensuring all samples met the required standards. The reads were aligned to the rat reference genome (rn7) using HISAT2, and feature quantification was achieved with feature counts.

Downstream analyses, including normalization and differential expression analysis, were conducted using limma-voom. Differentially expressed genes (DEGs) were identified using the following criteria: a *p*-value threshold of 0.05 and absolute fold change cutoff of 1. Gene Ontology Enrichment Analysis was performed using SRplot (Tang et al., 2023). Finally, Gene Set Enrichment Analysis was performed using the GSEA software (Broad Institute) to identify enriched gene sets. Expression values of the selected genes were standardized to z-scores and visualized in a heatmap. For each gene, the z-score was calculated as follows: Z=(X−μ) /σ, where X is the expression value in an individual sample, μ is the mean expression across the control group, and σ is the standard deviation of expression values within that control group.

### 2.10 RT-qPCR

RNA isolated from the ipsilateral and contralateral dorsal horn of the spinal cord was reverse transcribed into complementary DNA (cDNA) using a high-capacity cDNA synthesis kit (Applied Biosystems, Warrington, UK). For quantitative RT-PCR (RT-qPCR), each reaction well contained 1 μl of cDNA 5 μl of SYBR Green (Applied biosystems, Warrington, UK), 3.5 μl of RNase-free H_2_O, and 0.5 μl of gene-specific primers (a mix of forward and reverse primers at a concentration of 10 μM). Each sample was run in triplicate, and the results were normalized to the expression of the housekeeping genes *β-actin* and *gapdh*, expressed as a percentage of the control group (no RRS + sham) and calculated using the ΔΔCT method. Primer sequences are listed in Table 1.

**Table 1:**
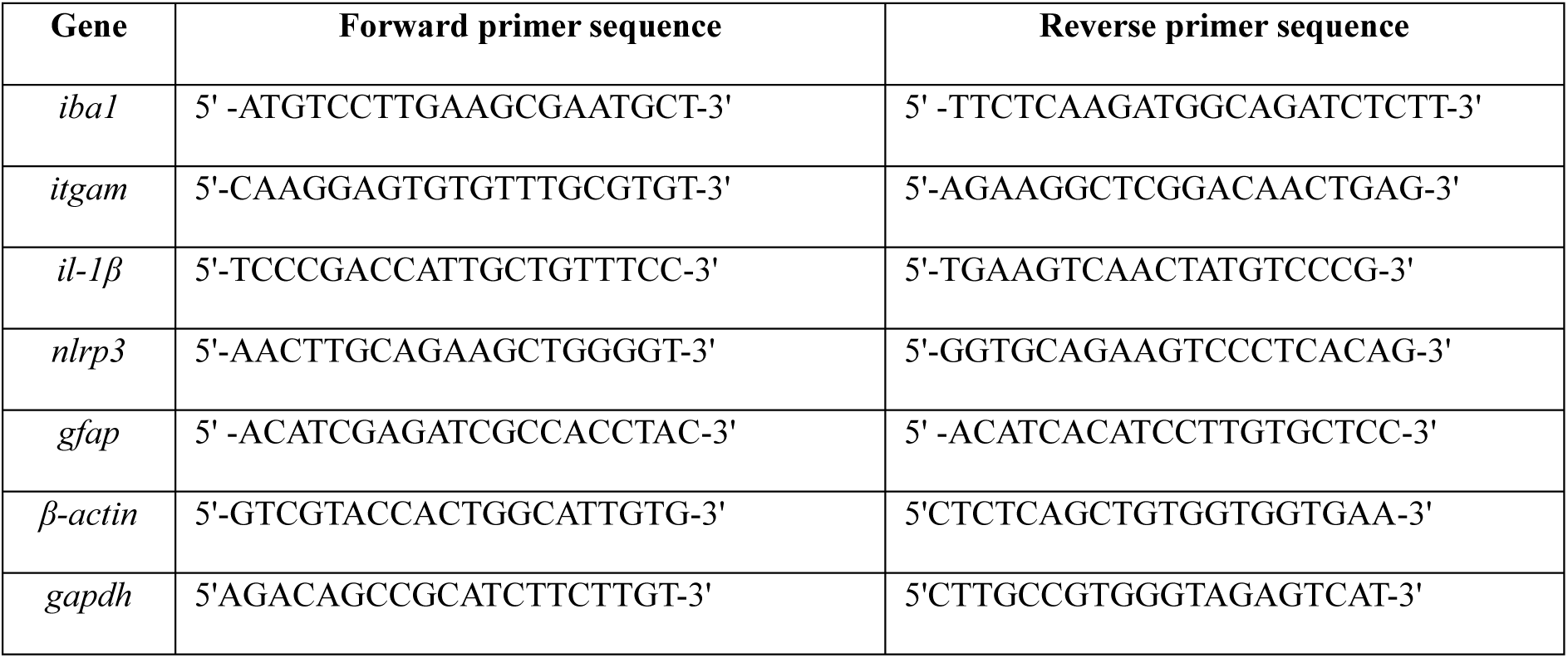
Primer sequences.

### 2.11 Immunohistochemistry for Iba1

Rats were transcardially perfused with phosphate buffer (PB) and 4% paraformaldehyde (PFA) in PB, and the spinal cords were post-fixed overnight at 4 °C followed by storage in 20% sucrose. The lumbar spinal cord segments (L4–L6) were sectioned at 30 µm, blocked in 5% goat serum, incubated overnight with rabbit anti-Iba1 (1:1000, WAKO, 019-19741), washed and incubated with secondary antibody conjugated to Alexa Fluor 594 (1:500).

Fluorescent images were acquired using a Zeiss Axio Imager 2 microscope. Maximum intensity projections of z-stacks and tile scans were generated using Zen Blue software (ZEISS Microscopy Software) to visualize and analyse high-resolution image datasets. Image acquisition settings, including objective magnification (40x), exposure time (100 ms), and z-stack step size (2 µm), were kept consistent across samples. For subsequent analyses, 3-4 sections per animal were used. Iba1-positive cells were quantified using Imaris 10.1 (Oxford Instruments). Fluorescence intensity-based analyses were performed in ImageJ (Fiji) by measuring mean pixel intensity in manually defined regions of interest (ROIs) within the ipsilateral and contralateral dorsal horn. Background subtraction and normalization methods were applied to ensure accurate quantification.

### 2.12 Statistical analysis

Data was analysed using GraphPad Prism, version 9.0 (GraphPad Software, Inc., La Jolla, CA, USA), which was also used to generate graphs. Prior to statistical testing, all datasets were assessed for normality and homogeneity. Comparisons between two groups, such as for FST and faecal corticosterone data, were performed using unpaired student’s t-test. Data from von Frey and Hargreaves’ tests were examined using two-way repeated measures ANOVA followed by Newman-Keuls post-hoc test where appropriate. Behavioural tests including the PEAP, OFT, EPM, LDB and emotionality z-scores were evaluated with two-way ANOVA followed by Newman-Keuls post-hoc test where appropriate. Changes in gene expression (qPCR), microglia numbers and fluorescence intensity in the dorsal horn were analysed using three-way ANOVA followed by Newman-Keuls post-hoc test. For clarity, the results of the ANOVA comparisons are presented in Supplementary Table S1. Statistical significance was set at *p* < 0.05, and results are expressed as group means ± standard error of the mean (SEM).

## 3. RESULTS

### 3.1 RRS elicits a depressive-like phenotype and increases the magnitude and duration of mechanical and heat hypersensitivity post-surgery

Data revealed that while all RRS protocols elicit a decrease in body weight gain (Figure S2) only RRS for 6h/day for 21 days resulted in a depressive-like phenotype and increased the magnitude and duration of post-surgical nociceptive responding. Data pertaining to the other RRS protocols are presented in Figures S3 and S4. RRS (6h/day for 21days) significantly increased immobility time (*t*(22)=3.666, *p*<0.01) in the FST (Figure 1a) and faecal corticosterone levels were significantly elevated compared to non-RRS counterparts (*t*(28)=2.847, *p*<0.01) (Figure 1b), confirming an altered stress response and depressive-like phenotype of animals subjected to this RRS protocol.

**Figure 1:**
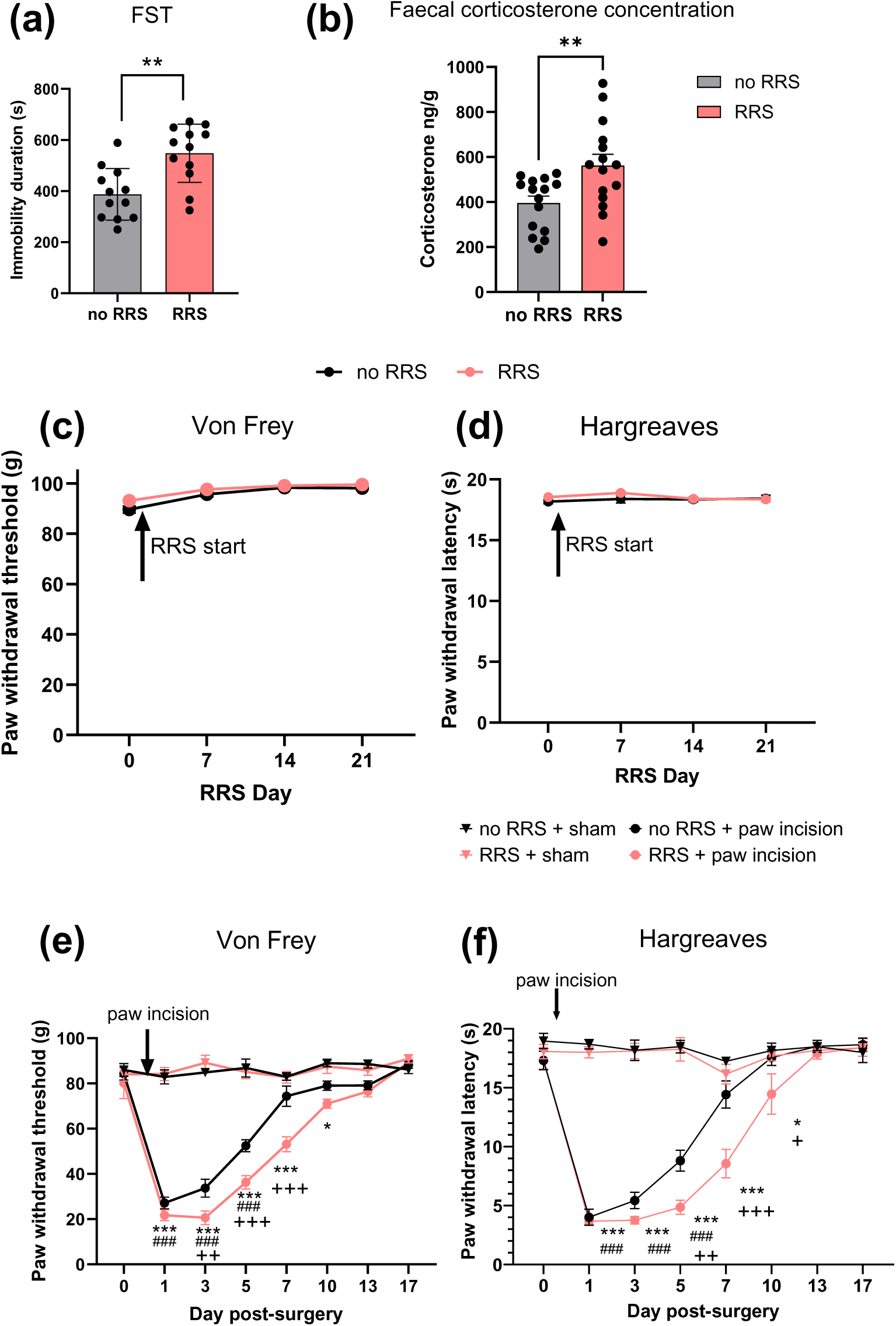
The effect of RRS on (a) immobility time in the FST (\*\**p*< 0.01) (b) faecal corticosterone levels (\*\**p*< 0.01), (c) mechanical and (d) thermal heat sensitivity, n=12-15/group. The effect of RRS on (d) paw withdrawal threshold (von Frey) and (e) paw withdrawal latency (Hargreaves test) post paw-incision or sham surgery. ^###^*p*<0.001 no RRS + sham vs. no RRS + paw incision, \**p*<0.05, \*\*\**p*<0.001 RRS + sham vs. RRS + paw incision, ^+^*p*<0.05, ^++^*p*<0.01 ^+++^*p*<0.001 no RRS + paw incision vs RRS + paw incision. n=6/group.

Analysis revealed that RRS did not alter paw withdrawal thresholds (PWTs) (Figure 1c) or paw withdrawal latencies (PWLs) (Figure 1d) at any point throughout the RRS period (days 0, 7, 14, 21). However, following paw incision or sham surgery, PWTs and PWLs were significantly reduced in rats that received paw incision compared to sham groups, indicating the development of post-surgical mechanical and heat hypersensitivity (Figure 1e-f). RRS exacerbated post-surgical mechanical hypersensitivity on days 3, 5 and 7 and prolonged mechanical hypersensitivity to day 10 post-surgery compared to non-RRS counterparts, which returned to baseline by day 7 (Figure 1e). RRS also exacerbated post-surgical heat hypersensitivity on days 5 and 7 post-surgery (Figure 1f) and prolonged heat hypersensitivity to day 10 compared to non-RRS counterparts. Taken together, these results demonstrate that RRS exacerbates and prolongs mechanical and heat hypersensitivity post-surgery.

### 3.2 RRS increases pain-related aversion and anxiety-like responding post-surgery

Assessment of PWTs revealed that paw incision surgery induced mechanical hypersensitivity from day 1 post surgery, an effect exacerbated and prolonged in animals pre-exposed to RRS (no RRS + paw incision vs RRS + paw incision) (Figure 2a), reconfirming the establishment of the model. To evaluate the emotional component of post-surgical pain and investigate whether RRS alters this state, a series of behavioural tests was conducted for up to 10 days post-surgery. On Day 2 post-surgery, we showed that animals that underwent paw incision spent increased time in the aversive bright compartment of the PEAP test, and this effect further increased in animals subjected to prior RRS (Figure 2b). In the OFT (day 4 post-surgery), animals subjected to RRS (no surgery) spent less time in the central zone compared to non-stressed counterparts (Figure 2c). Similarly, animals that underwent surgery, whether exposed to RRS or not, also spent less time in the central zone of the OFT. In the EPM (day 6 post-surgery), RRS or paw incision alone resulted in a decrease in the time spent in the open arms, but their combination did not further modify this response (Figure 2d). On day 10 post-surgery, animals exposed to RRS + paw incision spent more time in the bright compartment of the LDB, compared to rats that underwent RRS or surgery alone (Figure 2e). Z-scores were calculated to examine overall effect on emotionality and revealed that RRS and paw incision alone increase emotionality, an effect further heightened in animals subjected to RRS + surgery (Figure 2f). Together, these findings demonstrate that RRS exacerbates both the sensory and the affective components of post-surgical pain and amplifies pain-related aversion.

**Figure 2:**
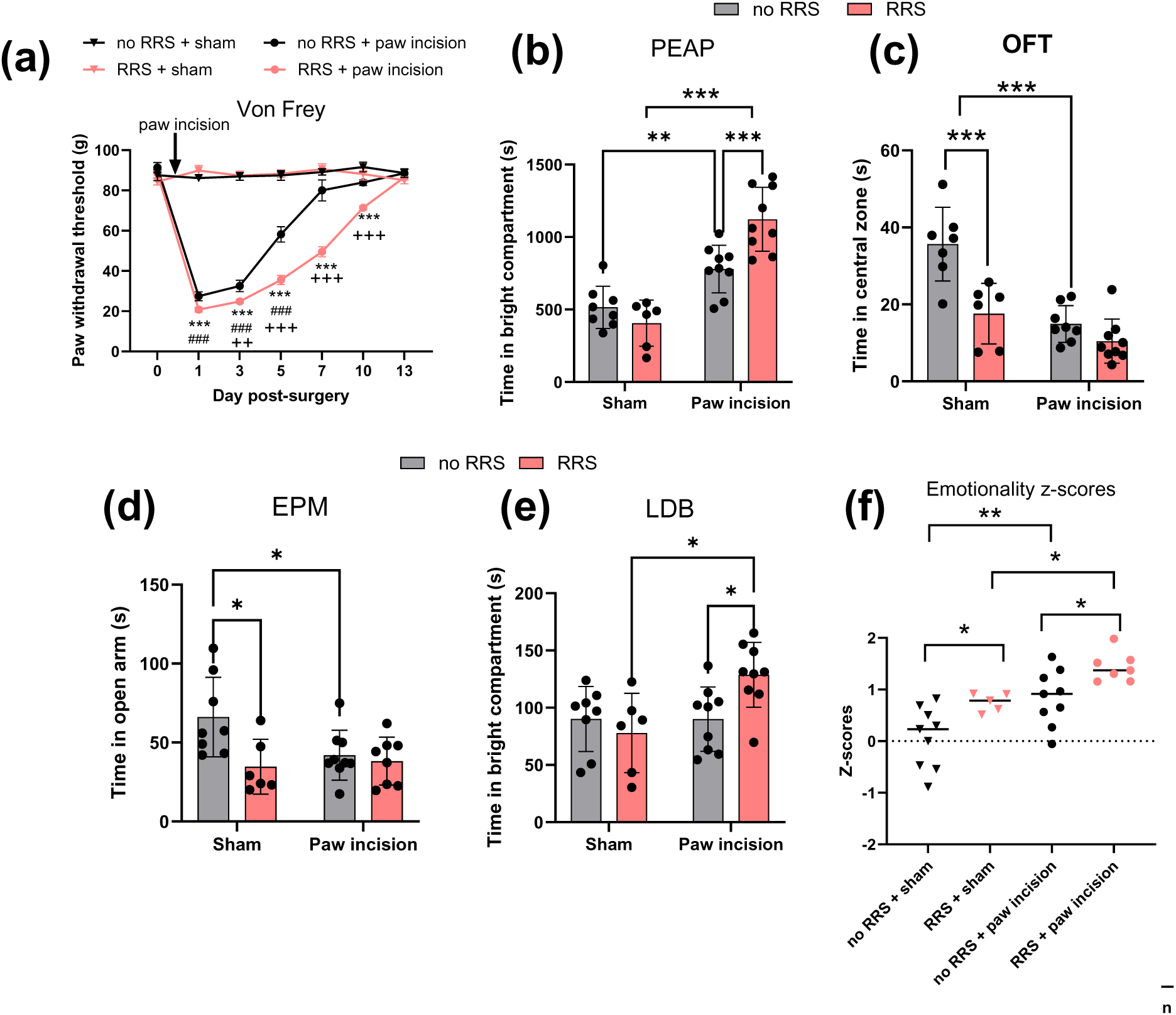
The effect of RRS on (a) paw withdrawal threshold (von Frey) ^###^*p*<0.001 no RRS + sham vs. no RRS + paw incision, \*\*\**p*<0.001 RRS + sham vs. RRS + paw incision, ^++^*p*<0.01 ^+++^*p*<0.001 no RRS + paw incision vs RRS + paw incision. (b) pain-related aversion (PEAP) and anxiety like behaviour in the (c) open field test (OFT), (d) elevated plus maze (EPM) and (e) light dark box (LDB). (f) overall emotionality (z-score). n=6-8 per group (\**p* < 0.05, *** *p* < 0.001, *** *p* < 0. 001).

### 3.3 RRS induces transcriptional alterations in the spinal NLRP3-IL-1β pathway post-surgery

To examine the molecular mechanisms underlying RRS-induced alterations in post-surgical pain, we performed RNA sequencing to examine gene expression changes in the ipsilateral dorsal horn five days post-surgery. This time point was chosen as it marks the emergence of significant differences in sensory hypersensitivity between surgery animals exposed, or not exposed, to chronic stress (no RRS + paw incision, RRS + paw incision, see Supplementary Figure 4e-f for behavioural analysis). At a nominal *p*-value threshold (*p* < 0.05), 946 genes were found to be significantly differentially expressed in the RRS + paw incision group compared to no RRS + paw incision controls (Figure 3a). Gene Ontology (GO) analysis highlighted significant enrichment of terms related to wound healing (*tnf, p2rx1, wnt4, c1qtnf1* etc), gliogenesis and glial cell differentiation *(nfib, bmerrb1, hexb, nrros, ctnnb1* etc) (Figure 3b), suggesting a potential role of neuroimmune mechanisms in the observed phenotype. To further investigate this, we performed Gene Set Enrichment Analysis (GSEA), to interrogate a well-characterized list of 42 genes involved in inflammasome-mediated signalling. Among these, 11 genes were found to contribute to a significant positive enrichment of the inflammasome-mediated signalling pathway (normalized enrichment score = 1.69, p < 0.05, FDR < 0.1) in RRS + paw incision rats (Figure 3c). Notable upregulated genes in the RRS + paw incision vs no RRS + paw incision group included *nlrp3*, *tlr4*, *p2rx1*, and other genes mediating activation of inflammasome signalling (Figure 3d). Further analysis on the effects of RRS, paw incision and their combination on the transcriptomic profile of the spinal cord are presented in Figures S5, S6 and S7.

**Figure 3:**
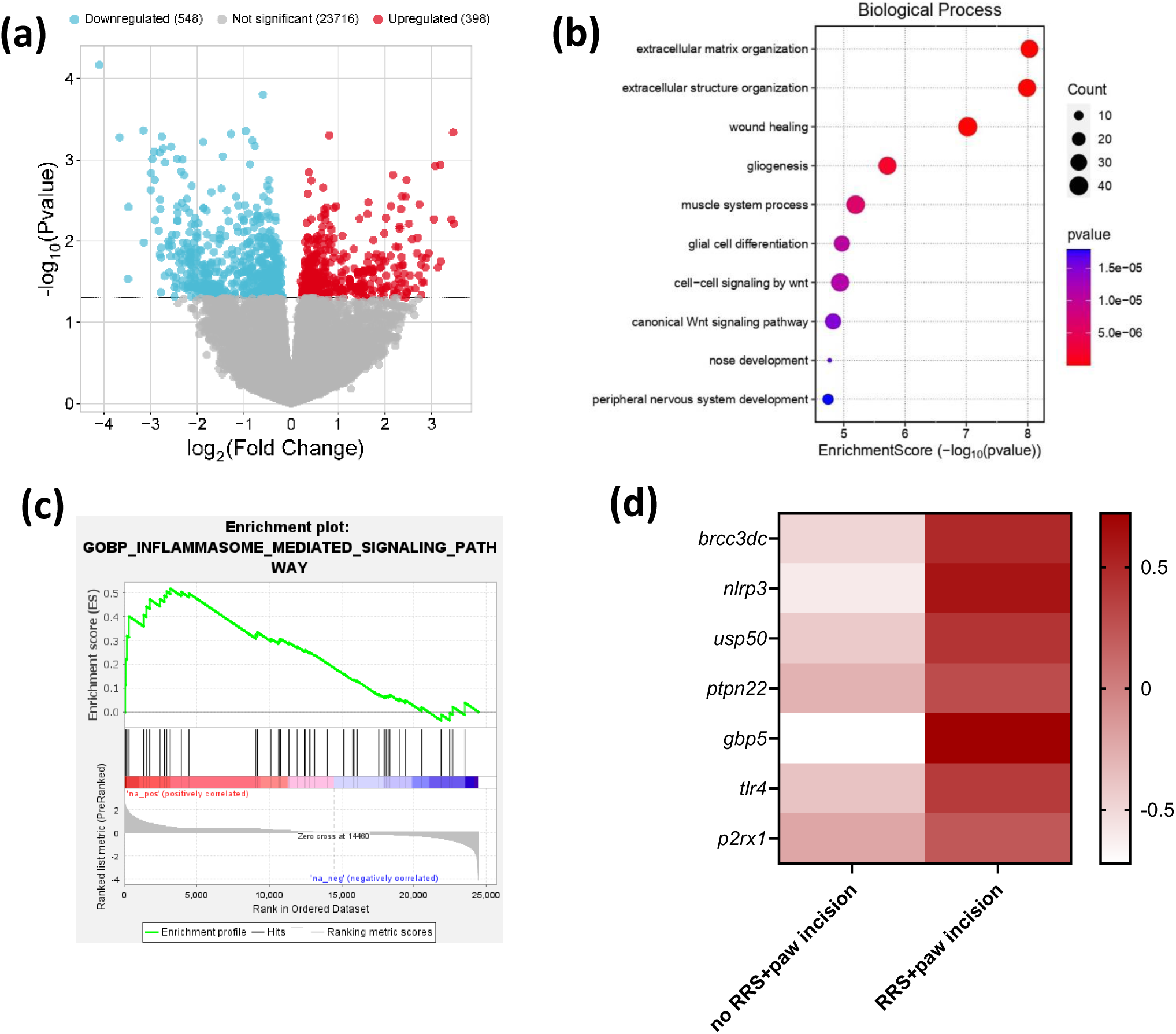
(a) Volcano plot depicting the 946 genes differentially expressed (nominal *P* values<0.05) between no RRS + paw incision (*n* = 4) and RRS + paw incision (*n* = 4) groups. (b) Top biological processes in the Gene Ontology pathway enrichment analyses of differentially expressed genes. (c) Gene set enrichment analysis (GSEA) significantly enriched gene set, *p* value = 0.005. (d) Upregulation of NLRP3 inflammasome pathway-related genes *(brcc3dc*, *P* = 0.094;*nlrp3*, *P* = 0.071; *usp50*, *P* = 0.163; *ptpn22*, *P* = 0.507; *gbp5*, *P* = 0.074; *tlr4*, *P* = 0.43; *p2rx1*, *P* = 0.027. Z-scores were calculated for each gene across all samples to standardize relative expression levels.

To validate the RNA sequencing data, RT-qPCR analysis was carried out on the ipsilateral and contralateral dorsal horn. RRS + paw incision was associated with significant upregulation of the expression of microglial markers *iba1* and *itgam* (Figure 4a-b,), as well as *nlrp3* and *il-1β* (Figure 4d-e), but not *gfap* (Figure 4c), compared to their non-stressed counterparts. Immunohistochemical analysis of Iba1 staining in the dorsal horn of the spinal cord confirmed an overall effect of RRS (*F*_(1,84)_= 5.401, *p*=0.0225) on the number of Iba1-positive cells, although *post hoc* analysis failed to reveal a significant difference between the groups (Figure 4g). However, analysis of fluorescence intensity revealed an increase in Iba1 levels in the ipsilateral dorsal horn of RRS + paw incision group compared to the contralateral side and to animals exposed to RRS alone (Figure 4h). Together, these findings indicate enhanced microglial activation and the upregulation of neuroinflammatory pathways, with a particular emphasis on the NLRP3-IL-1β inflammasome pathway, as potential contributors to the exacerbation of post-surgical pain in RRS rats.

**Figure 4:**
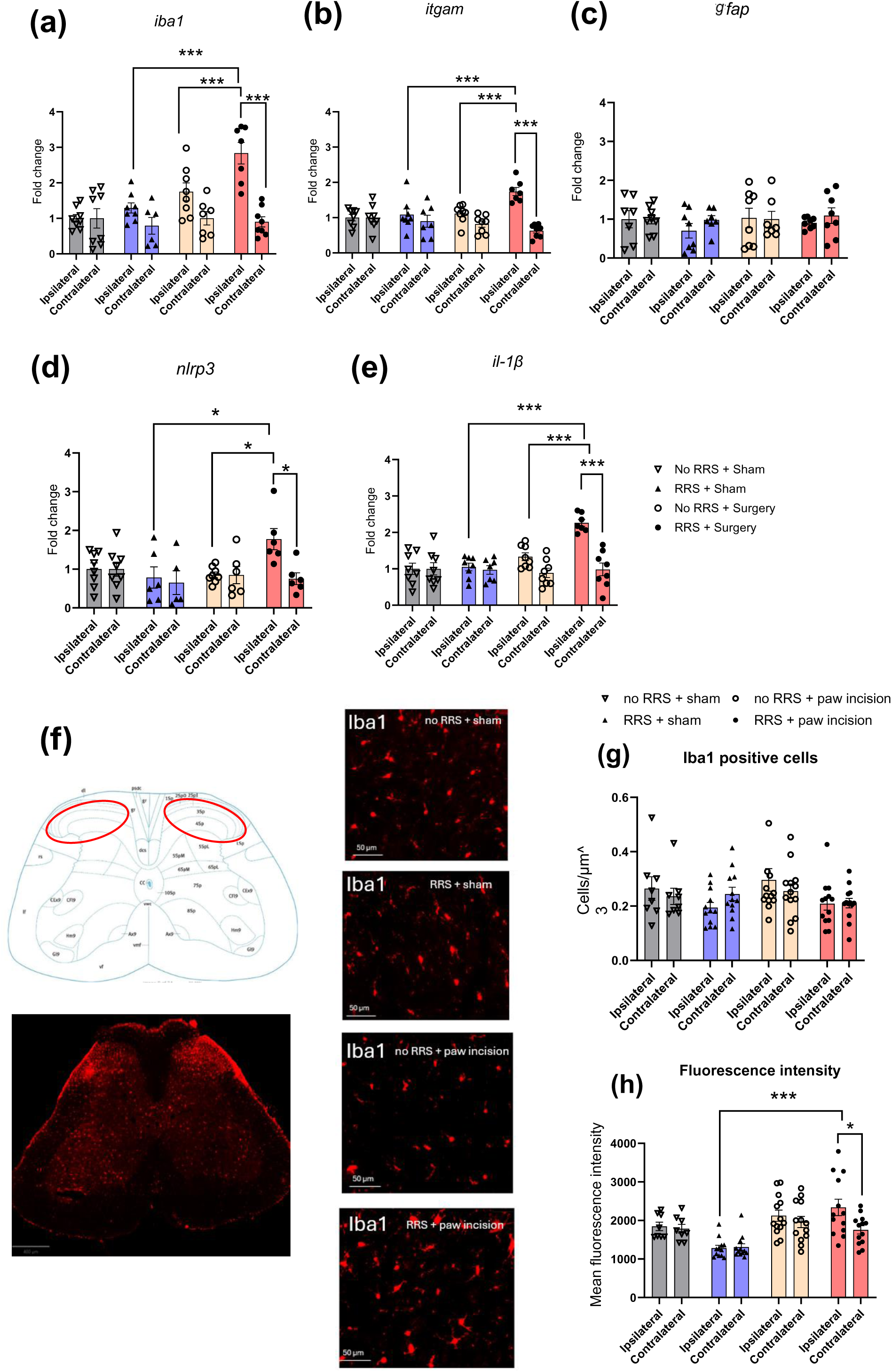
Relative gene expression of (a) *iba1*, (b) *itgam*, (c) *gfap*, (d) *nlrp3* and (e) *il-1β* in the ipsilateral dorsal horn, n=5-8 per group. Data are normalized to housekeeping genes β*-actin* and *gapdh* and presented as mean ± SEM. (f) Representative photomicrographs of Iba1 positive staining (red) in the spinal cord and high magnification of Iba1 in the ipsilateral dorsal horn for each of the 4 groups: no RRS + sham (top left), RRS + sham (top right), no RRS + paw incision (bottom left) and RRS + paw incision (bottom right). (g) The number of Iba1-positive cells and (h) mean fluorescence intensity of Iba1 staining in the region of interest (red circle) from ipsilateral and contralateral dorsal horn. \**p*<0.05, \*\*\**p*<0.001.

### 3.4 Blockade of spinal NLRP3 or IL-1β signaling attenuates RRS-induced exacerbation of pain-related aversion and mechanical hypersensitivity post-surgery

After identifying the spinal NLRP3-IL-1β pathway as a possible mediator of the RRS-induced enhancement of post-surgical pain, we then proposed to examine the role of this pathway in the behavioural effects observed using a pharmacological approach. As previously observed, pre-exposure to RRS increased the time spent in the bright compartment of the PEAP on day 2 post-surgery (Figure 5a and 5c,). Intrathecal administration of the NLRP3 inhibitor MCC950 (Figure 5a) or IL-1Ra (Figure 5c) attenuated the RRS-induced increase in time spent in the bright chamber of the PEAP chamber. These results suggest that inhibition of the spinal NLRP3-IL-1β pathway prevents the RRS-induced increase in the affective component of post-surgical pain.

**Figure 5:**
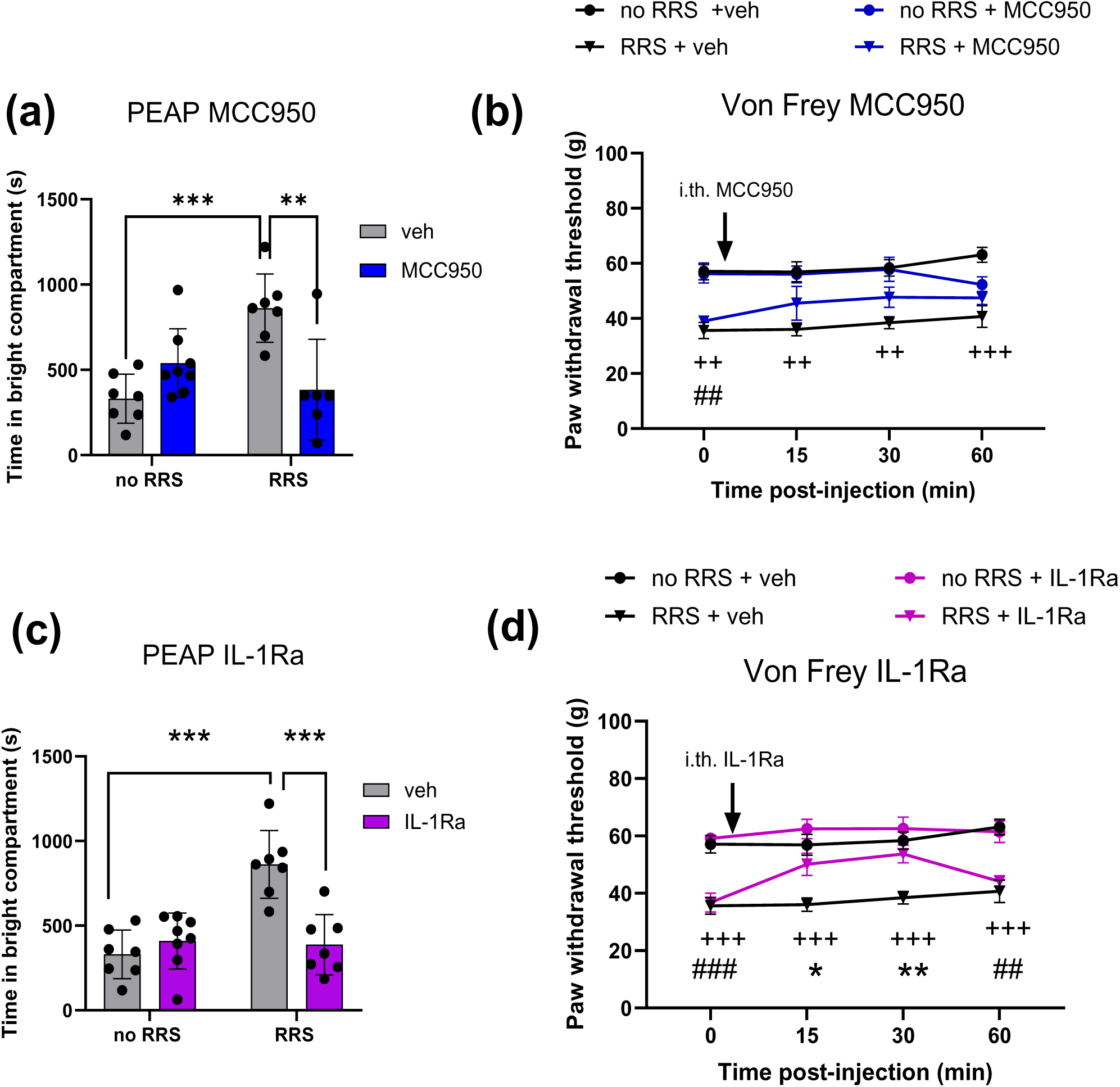
The effect of intrathecal administration of MCC950 and IL-1Ra on (a&c) pain-related aversion (PEAP; day 2 (**P<0.01 ***P<0.001) and (b&d) paw withdrawal thresholds (von frey; Day 5) following paw incision surgery. ^++^*p*<0.01, ^+++^*p*<0.001 No RRS + veh vs RRS + veh, ^##^*p*<0.01, ^###^*p*<0.001 no RRS + IL-1Ra/MCC950 vs RRS + IL-1Ra/MCC950, \**p*<0.05, \*\**p*<0.01 RRS + veh vs RRS + IL-1Ra. n=6-8 per group.

On day 5 post-surgery, our data re-confirmed that RRS exacerbates mechanical hypersensitivity (RRS + veh vs no RRS + Veh; Figure 5b and 5d, timepoint 0). Intrathecal administration of MCC950 (Figure 5b) or IL-1Ra (Figure 5d) did not alter paw withdrawal thresholds of non-stressed paw incision animals. However, IL-1Ra significantly attenuated surgery-induced mechanical hypersensitivity in animals pre-exposed to RRS compared to vehicle counterparts (RRS-Veh vs RRS-IL1Ra; 30 & 60 min). It was also noted that there was no significant difference between no RRS and RRS groups that received MCC950. These data suggest that increased spinal NLRP3-IL-1β signalling mediates RRS-induced exacerbation of mechanical hypersensitivity.

### 3.5 Chronic RU486 does not attenuate RRS-induced exacerbation of post-surgical hypersensitivity, pain-related aversion and anxiety behaviour

In order to determine if the effects of RRS on despair-like behaviour, post-surgical hypersensitivity, pain-related aversion and anxiety are mediated by glucocorticoid receptor signalling, rats received the glucocorticoid receptor antagonist RU486 prior to each RRS session. RU486 prevented the RRS-induced increase in immobility in the FST (Figure 6a), suggesting that blockade of glucocorticoid signalling can prevent RRS-induced increase in depressive-like behaviour.

**Figure 6:**
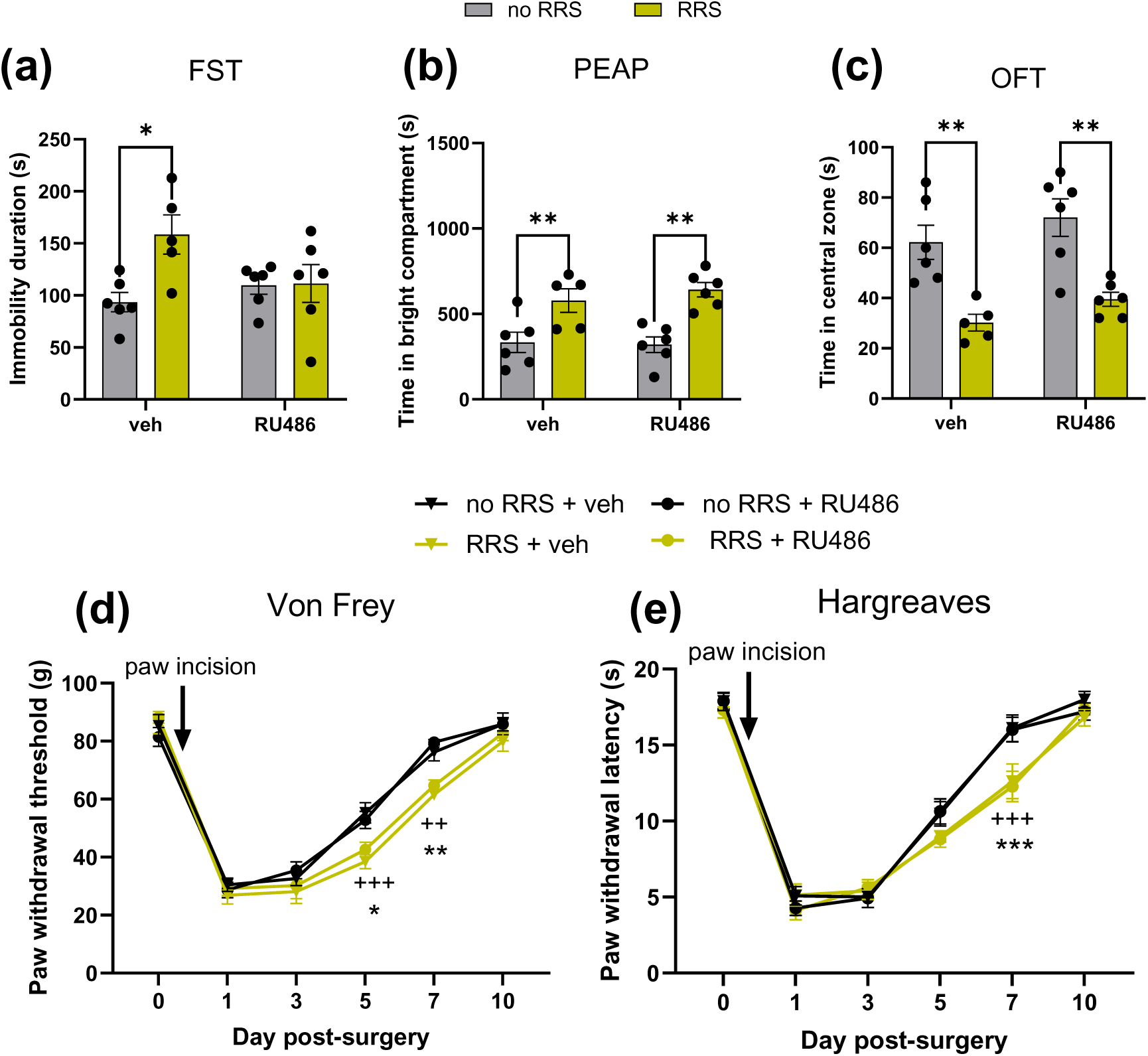
The effect of chronic administration of glucocorticoid antagonist RU486 (21 days) on (a) immobility in the FST (\**p* < 0.05), (b) pain-related aversion (PEAP Day 2 post surgery; **P<0.01) (c) time in the centre zone of the OFT (Day 4 post surgery, **P<0.01), (d) paw withdrawal threshold (von Frey) and (e) paw withdrawal latency (Hargreaves test). ^++^*p* <0.01, ^+++^*p* <0.001 no RRS + paw incision + veh vs RRS + paw incision + veh, * *p* < 0.05, ** *p* < 0.01, *** *p* < 0.001 no RRS + paw incision + RU486 vs RRS + paw incision + RU486. n= 5-6 per group.

Post-surgery, there was no effect of RU486 on PWTs (Figure 6d), PWLs (Figure 6e), time spent in the bright compartment of the PEAP (Figure 6b), and time spent in the central zone of the OFT (Figure 6c) in either RRS or no RRS exposed animals. These data suggest that RRS-induced glucocorticoid receptor signalling is not responsible for the exacerbated somatosensory or affective pain responding post-surgery.

### 3.6 Chronic propranolol attenuates RRS-induced exacerbation of post-surgical hypersensitivity, pain-related aversion and anxiety behaviour

To test whether the RRS-induced SNS and β-adrenergic receptor activation mediates the effects of RRS on despair-like behaviour, post-surgical hypersensitivity, pain-related aversion and anxiety, rats received the β-adrenergic antagonist propranolol prior to each RRS session. Propranolol prevented the RRS-induced increase in immobility in the FST (Figure 7a), suggesting that RRS-induced β-adrenergic activation contributes to the depressive-like phenotype observed.

**Figure 7:**
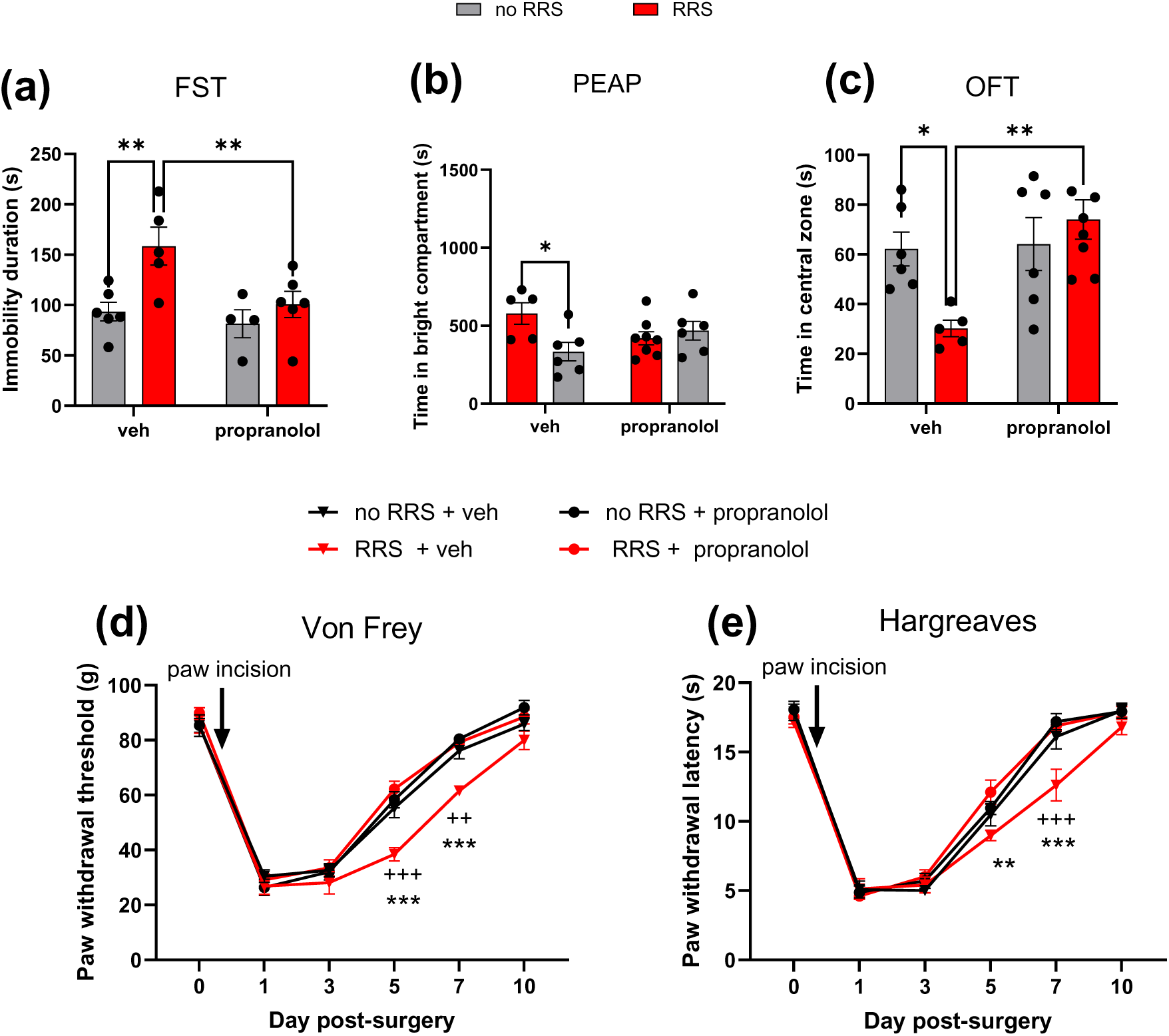
The effect of chronic administration of propranolol (21 days) on (a) immobility in the FST (\*\**p* < 0.01), (b) pain-related aversion (PEAP Day 2 post surgery; *P<0.05) (c) time in the centre zone of the OFT (Day 4 post surgery, *P<0.05 **P<0.01), (d) paw withdrawal threshold (von Frey) and (e) paw withdrawal latency (Hargreaves test). ^++^*p* <0.01, ^+++^*p* <0.001 no RRS + paw incision + veh vs RRS + paw incision + veh, * *p* < 0.05, ** *p* < 0.01, *** *p* < 0.001 RRS + paw incision + veh vs RRS + paw incision + propranolol. n= 5-8 per group.

Post-surgery, RRS-exposed rats that received propranolol exhibited increased paw withdrawal thresholds (Figure 7d) and paw withdrawal latencies (Figure 7e) compared to their vehicle-treated counterparts on days 5 and 7 (RRS + paw incision + veh vs RRS+ paw incision + propranolol). In addition, propranolol blocked the RRS-induced increase in the time spent in the bright compartment of the PEAP arena (Figure 7b) and the decrease in the time spent in the centre of the OFT arena (Figure 7c,) post-surgery. These data suggest that RRS-induced exacerbation of sensory and affective post-surgical pain is likely mediated by enhanced β-adrenergic activity.

## 4. DISCUSSION

Psychological factors such as chronic stress and depression are associated with increased and prolonged post-surgical pain, increased analgesic requirement and well recognised risk factors for the development of chronic post-surgical pain. This study demonstrated that chronic stress in the form of RRS results in altered stress responsivity and a depressive-like phenotype and exacerbates and prolongs sensory and affective pain responding post-surgery. Our data suggest that these behavioural effects are mediated by stress-induced β-adrenergic activity and enhanced spinal microglial activation and NLRP3/IL-1β signalling. These data provide further insight into the molecular and cellular mechanisms mediating the effects of chronic stress and negative affect on post-surgical pain.

We showed that repeated restraint stress (RRS) for 6h/day for 21 days induces both behavioural and physiological changes consistent with a depressive-like state. Specifically, we observed reduced body weight gain, increased immobility in the FST and elevated faecal corticosterone levels, mirroring prior studies that demonstrated similar effects following 21 days of restraint stress (6h/day) (Meng et al., 2021; Shoji & Miyakawa, 2020; Zhang et al., 2024). These results collectively underscore that prolonged RRS exposure not only elicits a depressive-like phenotype but also disrupts physiological homeostasis, highlighting a broader dysregulation of the stress response system.

Previous studies have reported that RRS alters mechanical withdrawal thresholds in the absence of injury. For example, RRS for 3 weeks (Imbe et al., 2012; Meng et al., 2021) in rats (6h/day) and 2 (Yang et al., 2024), 3 (Zhu et al., 2021) or 4 weeks (Borbely et al., 2023) in mice (6h/day) resulted in significant increases in mechanical and heat sensitivity, indicative of stress-induced hypersensitivity. In comparison, the data in the present study failed to find any effect of RRS on mechanical or heat sensory thresholds over the course of the 21 days of RRS. The discrepancy may stem from differences in the time of testing relative to stress exposure. For example, Imbe et al. measured mechanical sensitivity 1 hour after RRS exposure, while we assessed PWTs 18-20h post-RRS to capture the broader, cumulative effects of RRS sessions rather than any immediate post-stress responses.

Regarding the impact of RRS on post-surgical somatosensory responses, our findings indicate that RRS significantly heightened mechanical and heat hypersensitivity, particularly on post-surgical days 5 and 7, and prolonged hypersensitivity up to day 10. These results align with the published literature demonstrating that both acute and chronic pre-surgical stressors amplify the magnitude and prolong the duration of post-surgical somatosensory hypersensitivity in rodents (Bella et al., 2022). In particular our data align with that reported by, Meng and colleagues who demonstrated 21-day RRS protocol (6h/day) prolongs mechanical hypersensitivity up to day 10 following paw incision (Meng et al., 2021). However, to our knowledge, this is the first study to demonstrate that RRS also results in post-surgical heat hypersensitivity and enhanced affective pain responding. Notably, we found that paw-incised rats exhibited significant aversion to mechanical stimuli (PEAP test) as early as two days post-surgery. Comparable findings have been reported in a rat model of inguinal hernia repair, where animals demonstrated increased pain-related aversion within just three hours post-surgery (Bree et al., 2016). Interestingly, post-surgical pain-related aversion was further increased in rats subjected to prior RRS. These results suggest that RRS may increase unpleasantness and emotional distress, leading to an exaggerated avoidance of noxious stimuli compared to non-stressed paw-incised counterparts.

Regarding anxiety-like behaviour post-surgery, we observed that rats that underwent paw incision (without RRS) displayed anxiety-like behaviour on day 4 (OFT test) and day 6 post-surgery (EPM test) as well as an overall increase in emotionality score, data which align with previous studies showing that paw incision can induce anxiety-like behaviour in the OFT and EPM tests typically between 1 hour to 5 days post-surgery (Dai et al., 2011; Li et al., 2010). It has been suggested that anxiety-like behaviour could coincide with active hypersensitivity post-surgery, beginning to subside as pain thresholds return to baseline. Interestingly, prior RRS did not further enhance the anxiety-related behavioural phenotype observed post-surgery in the OFT and EPM, despite an exacerbation and prolongation of the somatosensory hypersensitivity. It is possible that this may be due to limitations of the behavioural tests, such as a potential floor effect, such that further increases in anxiety-related behaviour could not be detected. However, assessment of emotionality score across tests revealed a surgery-induced increase which was further exacerbated by prior RRS. Thus, taken together, prior chronic stress or a negative affective state is associated with enhanced and prolonged sensory and affective pain responding post-surgery.

It is well-documented that surgical procedures such as paw incision activate neuroinflammatory pathways, contributing to pain hypersensitivity (Arora et al., 2018; Beggs et al., 2012; Chen et al., 2018; Cui et al., 2025; Ito et al., 2009; Li et al., 2008; Liang et al., 2010; Romero-Sandoval et al., 2008; Sun et al., 2017; Wen et al., 2009; Zhao et al., 2025). Previous studies have shown that spinal microglial activation contributes to paw incision-induced hypersensitivity for up to 4 days post-surgery, coinciding with active hypersensitivity (Ito et al., 2009; Romero-Sandoval et al., 2008; Sun et al., 2017; Wen et al., 2009). Interestingly, in the current study, paw incision in the absence of prior RRS was not associated with any change in spinal microglial activity/density or the expression of proinflammatory markers 5 days post-surgery. Data suggest that as sensory hypersensitivity post-injury resolves, microglial activation is typically absent (Arora et al., 2018; Ito et al., 2009; Romero-Sandoval et al., 2008; Wen et al., 2009) and other glial and neuroimmune factors may come into play. Accordingly, on day 5 post-surgery, while animals still display mechanical and heat hypersensitivity, the magnitude of this response is reduced compared to earlier timepoints post-surgery. This is not the case for animals pre-exposed to RRS which continue to display robust sensory hypersensitivity and associated increases in microglial activation and proinflammatory mediator expression. This heightened neuroimmune and glial activity in animals pre-exposed to stress was further confirmed by transcriptomic analysis revealing alterations in inflammatory pathways, gliogenesis and glial cell differentiation.

Significant enrichment of the spinal NLRP3-IL-1β pathway in the ipsilateral dorsal horn of rats subjected to both RRS and paw incision was confirmed by RT-qPCR, highlighting this as a potential key mediator of the effects of RRS on post-surgical pain. Similarly, spinal microglial activation (Arora et al., 2018) accompanied by increased IL-1β (Sun et al., 2019; Sun et al., 2017) have been shown to underlie the effects of acute stress on post-surgical mechanical hypersensitivity and perioperative intrathecal inhibition of microglial activation decreased IL-1β expression and shortened the duration of SDS-induced post-surgical mechanical hypersensitivity (Sun et al., 2019). The data here expand and extend on these findings demonstrating that spinal NLRP3-IL1β signalling mediates RRS-induced potentiation of both somatosensory and affective aspects of post-surgical pain.

Glucocorticoid receptor inhibition effectively attenuates immobilization stress-induced prolongation of hypersensitivity to mechanical, heat, and cold stimuli in male rats (Cao et al., 2015). Similarly, glucocorticoid receptor inhibition mitigated the SPS-induced exacerbation of post-surgical mechanical hypersensitivity, microglial activation and cytokine expression (Wu et al., 2019). In contrast, the data herein revealed that while glucocorticoid receptor inhibition prevented the RRS-induced increase in despair-like behaviour, it did not prevent the exacerbation of post-surgical somatosensory hypersensitivity, aversion, or anxiety-like behaviour. These data suggest that pre-surgical despair-like behaviour is not a prerequisite for RRS-induced exacerbation of post-surgical somatosensory hypersensitivity, aversion, or anxiety-like behaviour. Taken together the data suggest that acute stress-induced exacerbation of post-surgical pain my involve glucocorticoid receptor signalling, while the effects of chronic stress may be mediated through alternative mechanisms.

We then sought to examine the role of SNS/β-adrenergic receptor activation on RRS-induced exacerbation and prolongation of post-surgical pain. Propranolol prevented the RRS-induced increase in despair-like behaviour, as well as the exacerbation of hypersensitivity, pain-related aversion and anxiety-like behaviour post-surgery. Our findings are in line with recent research suggesting that serum IL-6, induced by β3-adrenoceptor signalling in brown adipocytes, can lead to blood-spinal cord barrier disruption and spinal microglial activation, potentially contributing to acute stress-mediated persistent post-surgical hypersensitivity (Zhu et al., 2024). β-adrenergic receptors are known to induce microglial priming, and propranolol inhibits restraint stress-induced microglial activation (Sugama et al., 2019). Given that IL-6 and IL-1β are both key neuroimmune mediators involved in stress-induced microglial activation and pain modulation, we propose a mechanism in which chronic stress primes spinal microglia via β-adrenergic receptor signalling, and when combined with paw incision, this leads to upregulation of the NLRP3-IL-1β signalling, driving both sensory and affective pain exacerbation.

In conclusion, our findings demonstrate that RRS significantly exacerbates both somatosensory hypersensitivity and affective-pain dysregulation post-surgery, effects mediated, at least in part, by the β-adrenergic receptor activation, enhanced spinal microglial activation and NLRP3-IL-1β signalling. Together, these results provide valuable insights into how chronic stress and a negative affective state alters post-surgical pain processing and emotionality, suggesting potential targets for therapeutic interventions for the co-morbidity of mood and pain disorders.

## Acknowledgments

This work was supported by European Union’s Horizon 2020 research and innovation programme under the Marie Skłodowska-Curie grant agreement No 955684. This work was facilitated by the University of Galway, the Centre National de la Recherche Scientifique (contract UPR3212) and the University of Strasbourg. We would like to thank the Bio-Resources Unit and UMS3415 Chronobiotron for animal care and the In Vitro UAR 3156 imaging platform.

## Author contributions

Conceptualization MR, IY, DF; Formal analysis; AB, CD, DRA; Funding acquisition; MR, IY, DF; Investigation and Methodology AB, KA, CD, AMD, PH, CDM; Supervision; MR, IY, DF; Writing - original draft AB; Writing - review & editing AB, MR, IY, DF.

## Abbreviations

PPP: persistent post-surgical painy
RRS: repeated restraint stress
SPS: single prolonged stress
RSD: REM sleep disturbance
RDS: repeated defeat stress
PWT: paw withdrawal threshold
PWL: paw withdrawal latency
FST: forced swim test
PEAP: place escape/avoidance paradigm
OFT: open field test
EPM: elevated plus maze
LDB: light-dark box
BLA: basolateral amygdala
CeA: central amygdala
TNF-a: tumour necrosis factor a
IL-1β: interleukin 1β
NLRP-3: NLR family pyrin domain containing 3
Iba1: ionized calcium-binding adapter molecule 1
GFAP: glial fibrillary acidic protein
GR: glucocorticoid receptor
NF-κB: nuclear factor kappa-light-chain-enhancer of activated B cells
GAPDH: glyceraldehyde 3-phosphate dehydrogenase
DEGs: differentially expressed genes
GSEA: gene set enrichment analysis
IL-1Ra: interleukin 1 receptor antagonist
SNS: sympathetic nervous system

## Notes

### Competing Interest Statement

The authors have declared no competing interest.

