## Supplementary material for "Repeated restraint stress-induced increase in post-surgical somatosensory hypersensitivity and affective responding is mediated by β-adrenergic receptor activation and spinal NLRP3-IL1β signalling in male rats": Suppl data

Supplementary Figure S1: Experimental timeline for study 1 and 2 (created using Biorender.com)

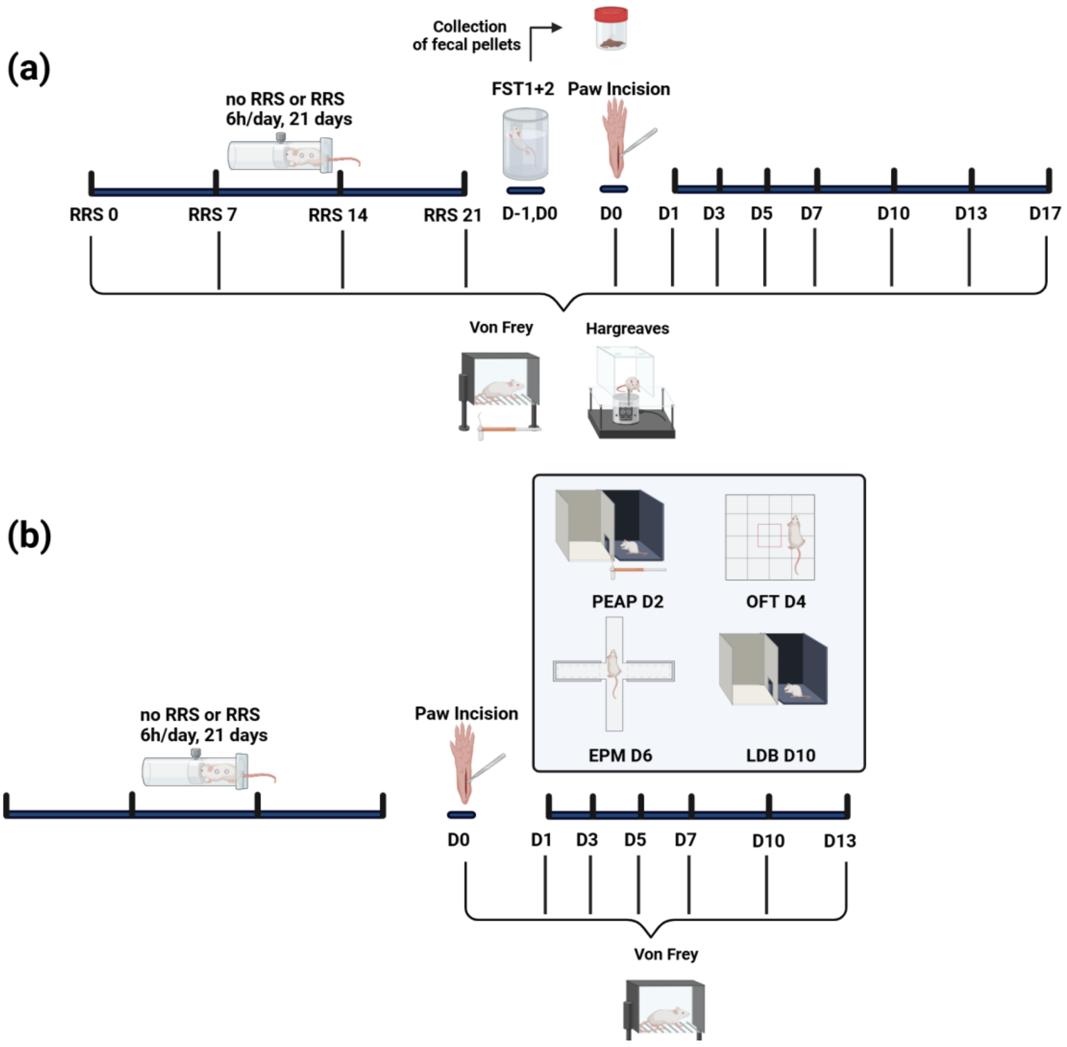

Supplementary Figure S2

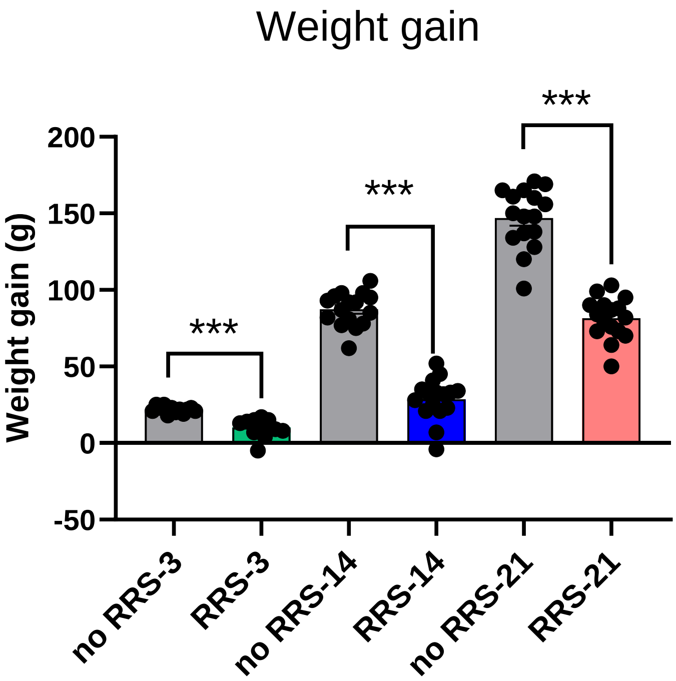


**Supplementary Figure 2:** The effect of RRS-3 (3 days, 6h/day), RRS-14 (14 days, 2.5h/day) and RRS-21 (21 days, 6h/day) on body weight gain. Data expressed as group means ± SEM, n=12-16/group. *** *p*< 0.001. no RRS vs RRS.

Supplementary Figure 3

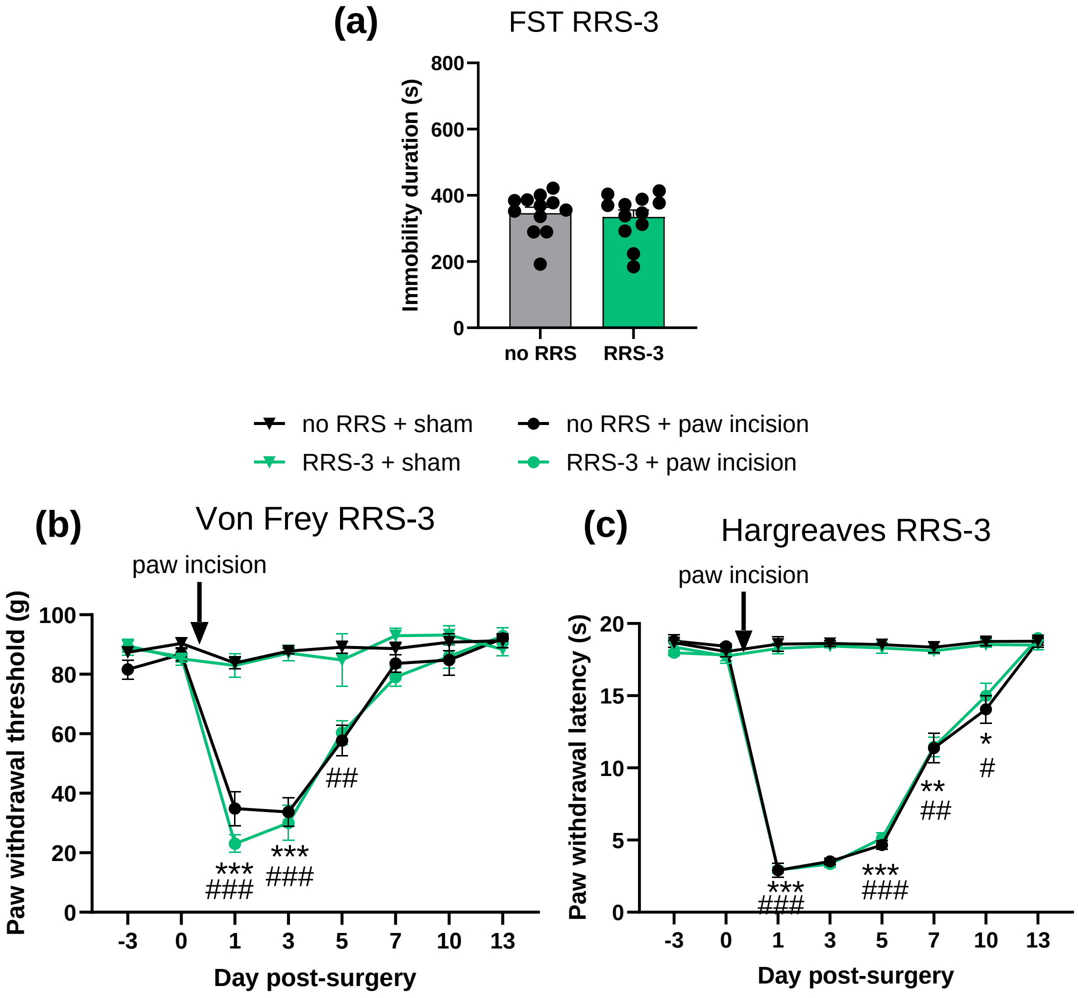


**Supplementary Figure 3:** The effect of RRS 6h/day for 3days (RRS-3) on (a) immobility time in the FST, (b) mechanical and (c) thermal heat hypersensitivity, n=6-12/group. Two-way repeated measures ANOVA followed by Newman-Keuls post-hoc: no RRS + sham vs no RRS + paw incision $^{\#}p<0.05$ , $^{\#\#}p<0.01$, $^{\#\#\#}p<0.001$ , RRS + sham vs RRS + paw incision $^{*}p<0.05$ , $^{**}p<0.01$, $^{***}p<0.001$. Data expressed as group means ± SEM.

Supplementary Figure 4

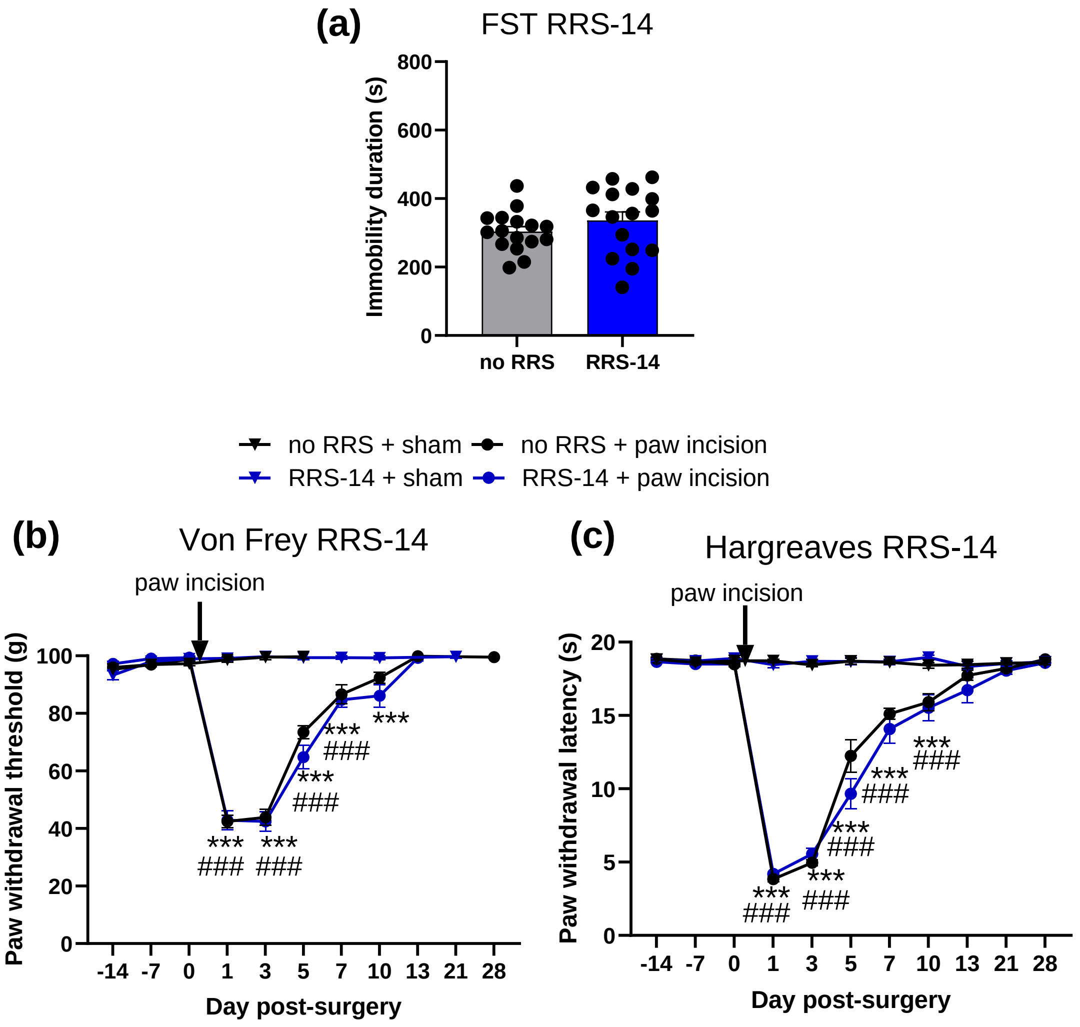


**Supplementary Figure 4:** The effect of RRS for 2.5h/day for 14 days (RRS-14) on (a) immobility time in the FST, (b) mechanical and (c) thermal heat hypersensitivity, n=8-16/group. Two-way repeated measures ANOVA followed by Newman-Keuls post-hoc: no RRS + sham vs no RRS + paw incision $^{###}p<0.001$ , RRS + sham vs RRS + paw incision $^{***}p<0.001$. Data expressed as group means ± SEM.

Supplementary Figure 5

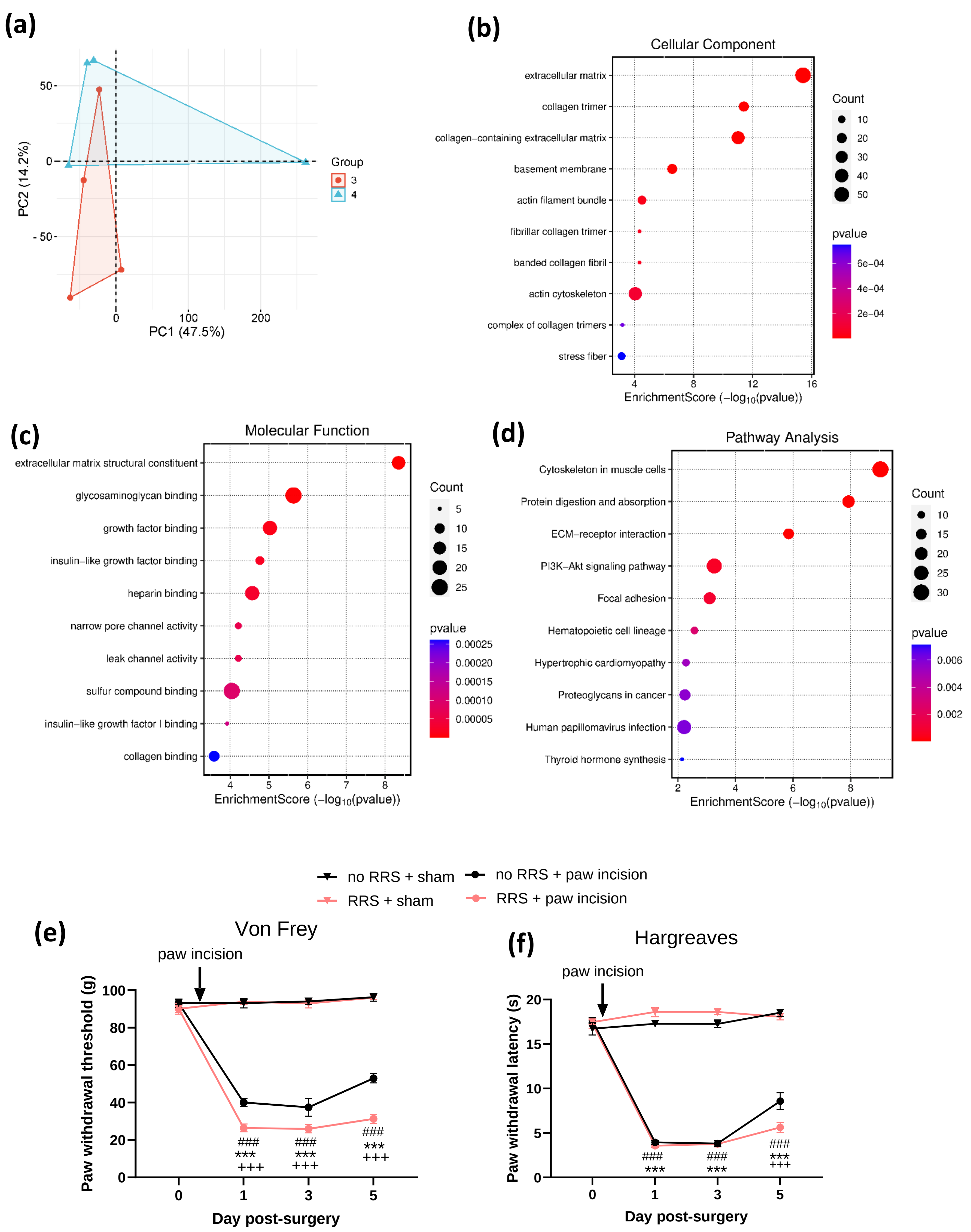

**Supplementary Figure 5:** (a) Principal component analysis between the groups no RRS + paw incision (n=4) and RRS + paw incision (n=4). GO enrichment analysis: (b) cellular component (c) molecular function and (d) KEGG pathways of the 946 differentially expressed genes (DEG, nominally significant) in animals exposed to RRS + paw incision. (e) Paw withdrawal thresholds and (f) paw withdrawal latencies of animals used for 3'RNAseq analysis post-surgery. $^{###}p<0.01$ no RRS + sham vs no RRS + paw incision, ***$p<0.001$ RRS + sham vs RRS + paw incision, $^{++}p<0.01$, $^{+++}p<0.001$. no RRS + paw incision vs RRS + paw incision. Data expressed as group means ± SEM, n=8/group.

Supplementary figure 6

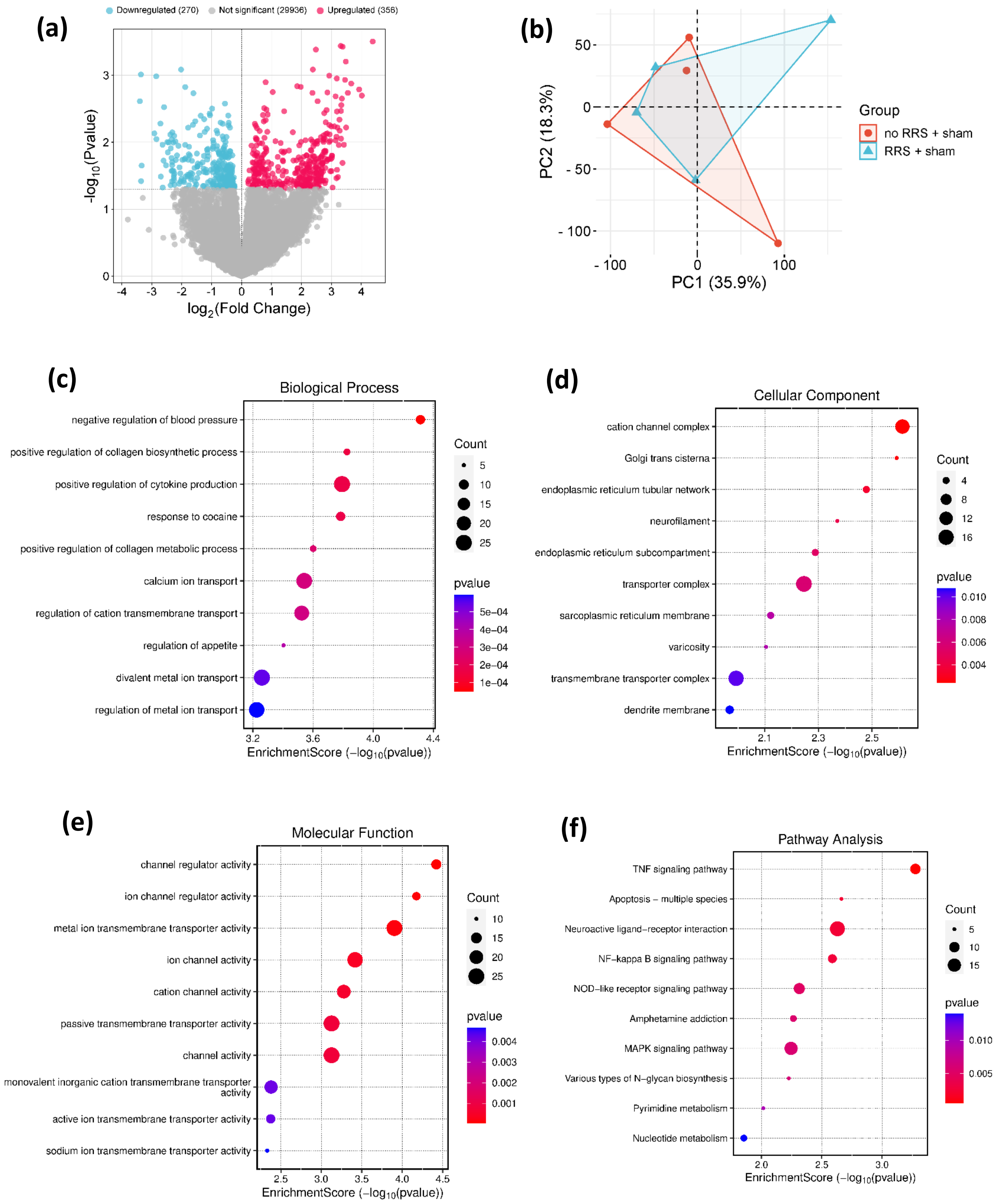

**Supplementary Figure 6:** (a) Volcano plot depicting the 628 genes differentially expressed (nominal *P* values<0.05) between no RRS + sham (*n* = 4) and RRS + sham (*n* = 4) groups. (b) Principal component analysis between the groups no RRS + sham and RRS + sham. GO enrichment analysis: (c) biological process, (d) cellular component, (e) molecular function and (f) KEGG pathways of the differentially expressed genes (DEG, nominally significant).

Supplementary figure 7

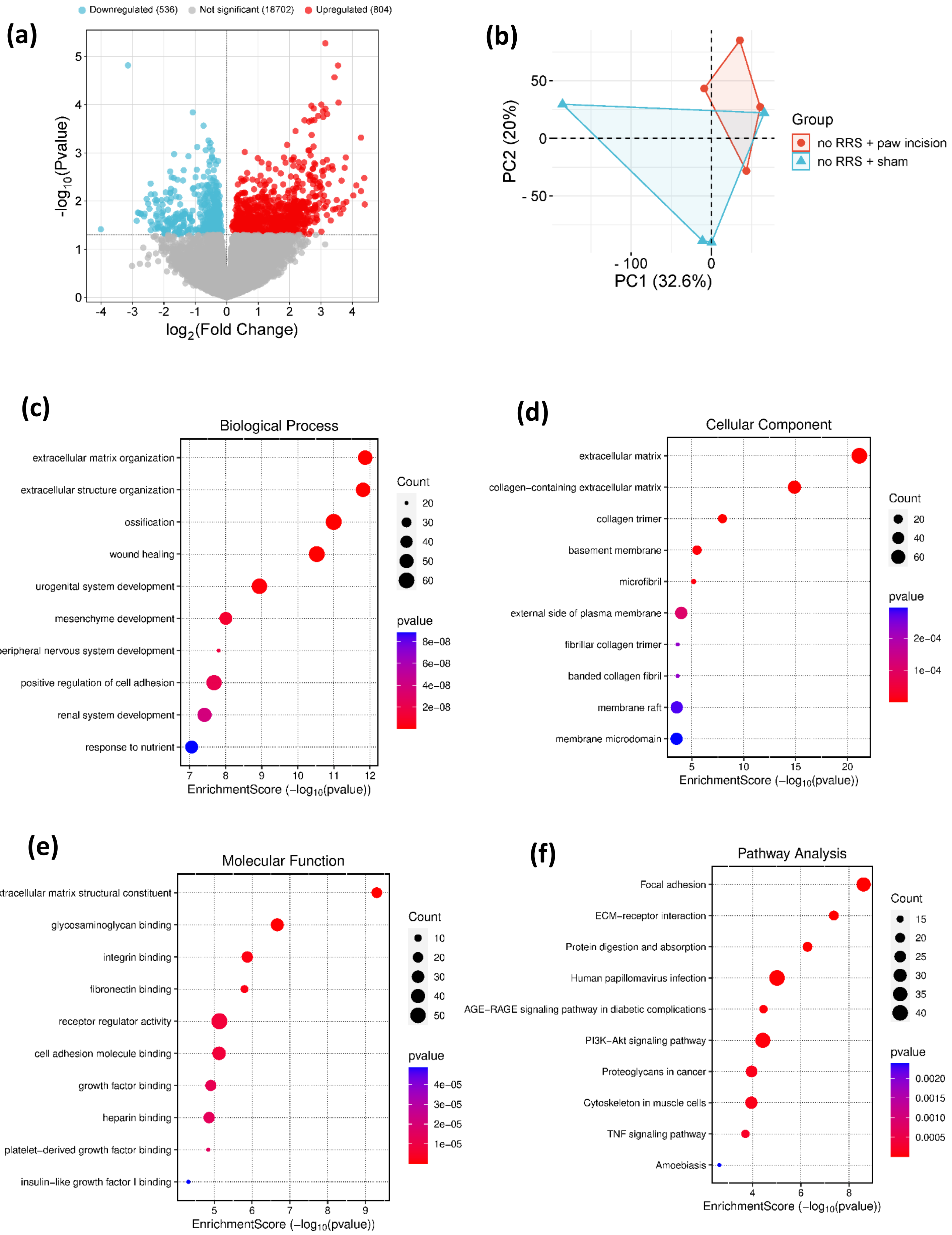

**Supplementary Figure 7:** (a) Volcano plot depicting the 1340 genes differentially expressed (nominal *P* values<0.05) between no RRS + sham (*n* = 4) and no RRS + paw incision (*n* = 4) groups. (b) Principal component analysis between the groups no RRS + sham and no RRS + paw incision. GO enrichment analysis: (c) biological process, (d) cellular component, (e) molecular function and (f) KEGG pathways of the differentially expressed genes (DEG, nominally significant).

**Supplementary Table 1:** Results of ANOVAs for data depicted in graphs. P in bold indicates significance.

| | ANOVA table | F (DFn, DFd) | P value |
|---|---|---|---|
| **Figure 1c** | Time X RRS | F (3, 66) = 0.8756 | P=0.4583 |
| | Time | F (3, 66) = 15.37 | **P<0.0001** |
| | RRS | F (1, 22) = 0.04881 | P=0.8272 |
| **Figure 1d** | Time X RRS | F (3, 66) = 0.1492 | P=0.9299 |
| | Time | F (3, 66) = 2.794 | **P=0.0471** |
| | RRS | F (1, 22) = 0.6571 | P=0.4263 |
| **Figure 1e** | Time | F (7, 140) = 81.26 | **P<0.0001** |
| | Paw incision | F (1, 20) = 610.8 | **P<0.0001** |
| | RRS | F (1, 20) = 16.83 | **P=0.0006** |
| | Time x Paw incision | F (7, 140) = 71.29 | **P<0.0001** |
| | Time x RRS | F (7, 140) = 2.073 | P=0.0503 |
| | Paw incision x RRS | F (1, 20) = 18.93 | **P=0.0003** |
| | Time x Paw incision x RRS | F (7, 140) = 1.732 | P=0.1061 |
| **Figure 1f** | Time | F (7, 140) = 83.43 | **P<0.0001** |
| | Paw incision | F (1, 20) = 296.3 | **P<0.0001** |
| | RRS | F (1, 20) = 11.88 | **P=0.0026** |
| | Time x Paw incision | F (7, 140) = 82.91 | **P<0.0001** |
| | Time x RRS | F (7, 140) = 2.900 | **P=0.0073** |
| | Paw incision x RRS | F (1, 20) = 5.000 | **P=0.0369** |
| | Time x Paw incision x RRS | F (7, 140) = 2.132 | **P=0.0440** |
| **Figure 2a** | Time | F (6, 156) = 129.1 | **P<0.0001** |
| | Paw incision | F (1, 26) = 953.3 | **P<0.0001** |
| | RRS | F (1, 26) = 49.01 | **P<0.0001** |
| | Time x Paw incision | F (6, 156) = 132.0 | **P<0.0001** |
| | Time x RRS | F (6, 156) = 4.301 | **P=0.0005** |
| | Paw incision x RRS | F (1, 26) = 40.68 | **P<0.0001** |
| | Time x Paw incision x RRS | F (6, 156) = 6.519 | **P<0.0001** |
| **Figure 2b** | Paw incision x RRS | F (1, 28) = 12.65 | **P=0.0014** |
| | Paw incision | F (1, 28) = 59.56 | **P<0.0001** |
| | RRS | F (1, 28) = 3.361 | P=0.0774 |
| **Figure 2c** | Paw incision x RRS | F (1, 26) = 6.878 | **P=0.0144** |
| | Paw incision | F (1, 26) = 28.89 | **P<0.0001** |
| | RRS | F (1, 26) = 18.90 | **P=0.0002** |
| **Figure 2d** | Paw incision x RRS | F (1, 27) = 4.120 | P=0.0523 |
| | Paw incision | F (1, 27) = 2.263 | P=0.1441 |
| | RRS | F (1, 27) = 6.648 | **P=0.0157** |
| **Figure 2e** | Paw incision x RRS | F (1, 28) = 5.816 | **P=0.0227** |
| | Paw incision | F (1, 28) = 5.737 | **P=0.0235** |
| | RRS | F (1, 28) = 1.560 | P=0.2220 |
| **Figure 2f** | Paw incision x RRS | F (1, 26) = 0.05132 | P=0.8226 |
| | Paw incision | F (1, 26) = 16.90 | **P=0.0003** |

| | | | |
|---|---|---|---|
| | RRS | F (1, 26) = 12.99 | **P=0.0013** |
| **Figure 4a** | Side | F (1, 52) = 27.81 | **P<0.0001** |
| | Paw incision | F (1, 52) = 15.87 | **P=0.0002** |
| | RRS | F (1, 52) = 3.146 | P=0.0820 |
| | Side x Paw incision | F (1, 52) = 13.02 | **P=0.0007** |
| | Side x RRS | F (1, 52) = 7.856 | **P=0.0071** |
| | Paw incision x RRS | F (1, 52) = 2.244 | P=0.1402 |
| | Side x Paw incision x RRS | F (1, 52) = 1.292 | P=0.2609 |
| **Figure 4b** | Side | F (1, 54) = 23.14 | **P<0.0001** |
| | Paw incision | F (1, 54) = 0.9069 | P=0.3452 |
| | RRS | F (1, 54) = 1.627 | P=0.2076 |
| | Side x Paw incision | F (1, 54) = 13.54 | **P=0.0005** |
| | Side x RRS | F (1, 54) = 9.153 | **P=0.0038** |
| | Paw incision x RRS | F (1, 54) = 1.866 | P=0.1776 |
| | Side x Paw incision x RRS | F (1, 54) = 3.589 | P=0.0635 |
| **Figure 4c** | Side | F (1, 53) = 0.7425 | P=0.3927 |
| | Paw incision | F (1, 53) = 0.4604 | P=0.5004 |
| | RRS | F (1, 53) = 0.4888 | P=0.4875 |
| | Side x Paw incision | F (1, 53) = 0.05895 | P=0.8091 |
| | Side x RRS | F (1, 53) = 0.9583 | P=0.3321 |
| | Paw incision x RRS | F (1, 53) = 0.3054 | P=0.5828 |
| | Side x Paw incision x RRS | F (1, 53) = 0.01577 | P=0.9005 |
| **Figure 4d** | Side | F (1, 45) = 4.005 | P=0.0514 |
| | Paw incision | F (1, 45) = 1.879 | P=0.1772 |
| | RRS | F (1, 45) = 0.2047 | P=0.6531 |
| | Side x Paw incision | F (1, 45) = 2.371 | P=0.1306 |
| | Side x RRS | F (1, 45) = 3.943 | P=0.0532 |
| | Paw incision x RRS | F (1, 45) = 5.771 | **P=0.0205** |
| | Side x Paw incision x RRS | F (1, 45) = 2.323 | P=0.1345 |
| **Figure 4e** | Side | F (1, 54) = 21.64 | **P<0.0001** |
| | Paw incision | F (1, 54) = 13.87 | **P=0.0005** |
| | RRS | F (1, 54) = 7.284 | **P=0.0093** |
| | Side x Paw incision | F (1, 54) = 17.93 | **P<0.0001** |
| | Side x RRS | F (1, 54) = 5.597 | **P=0.0216** |
| | Paw incision x RRS | F (1, 54) = 6.706 | **P=0.0123** |
| | Side x Paw incision x RRS | F (1, 54) = 3.796 | P=0.0566 |
| **Figure 4g** | Side | F (1, 84) = 0.04640 | P=0.8300 |
| | Paw incision | F (1, 84) = 0.1447 | P=0.7046 |
| | RRS | F (1, 84) = 5.401 | **P=0.0225** |
| | Side x Paw incision | F (1, 84) = 0.5410 | P=0.4641 |
| | Side x RRS | F (1, 84) = 2.180 | P=0.1436 |
| | Paw incision x RRS | F (1, 84) = 0.7059 | P=0.4032 |
| | Side x Paw incision x RRS | F (1, 84) = 0.1839 | P=0.6691 |
| **Figure 4h** | Side | F (1, 84) = 3.804 | P=0.0545 |
| | Paw incision | F (1, 84) = 24.02 | **P<0.0001** |

| | | | |
|---|---|---|---|
| | RRS | F (1, 84) = 6.654 | **P=0.0116** |
| | Side x Paw incision | F (1, 84) = 3.221 | P=0.0763 |
| | Side x RRS | F (1, 84) = 0.7116 | P=0.4013 |
| | Paw incision x RRS | F (1, 84) = 6.892 | **P=0.0103** |
| | Side x Paw incision x RRS | F (1, 84) = 1.617 | P=0.2070 |
| **Figure 5a** | RRS x Treatment | F (1, 24) = 18.11 | **P=0.0003** |
| | RRS | F (1, 24) = 5.393 | **P=0.0290** |
| | Treatment | F (1, 24) = 2.791 | P=0.1078 |
| **Figure 5b** | Time | F (3, 75) = 1.565 | P=0.2049 |
| | Treatment | F (1, 25) = 0.7684 | P=0.3891 |
| | RRS | F (1, 25) = 49.97 | **P<0.0001** |
| | Time x Treatment | F (3, 75) = 1.081 | P=0.3622 |
| | Time x RRS | F (3, 75) = 0.6920 | P=0.5598 |
| | Treatment x RRS | F (1, 25) = 5.413 | **P=0.0284** |
| | Time x Treatment x RRS | F (3, 75) = 0.8762 | P=0.4574 |
| **Figure 5c** | RRS x Treatment | F (1, 25) = 18.53 | **P=0.0002** |
| | RRS | F (1, 25) = 15.72 | **P=0.0005** |
| | Treatment | F (1, 25) = 9.488 | **P=0.0050** |
| **Figure 5d** | Time | F (3, 75) = 3.161 | **P=0.0295** |
| | Treatment | F (1, 25) = 9.916 | **P=0.0042** |
| | RRS | F (1, 25) = 108.7 | **P<0.0001** |
| | Time x Treatment | F (3, 75) = 2.621 | P=0.0569 |
| | Time x RRS | F (3, 75) = 1.217 | P=0.3096 |
| | Treatment x RRS | F (1, 25) = 2.888 | P=0.1016 |
| | Time x Treatment x RRS | F (3, 75) = 0.7364 | P=0.5336 |
| **Figure 6a** | RRS x Treatment | F (1, 19) = 4.985 | **P=0.0378** |
| | Treatment | F (1, 19) = 1.183 | P=0.2904 |
| | RRS | F (1, 19) = 5.542 | **P=0.0295** |
| **Figure 6b** | RRS x Treatment | F (1, 19) = 0.5072 | P=0.4850 |
| | Treatment | F (1, 19) = 0.2166 | P=0.6469 |
| | RRS | F (1, 19) = 27.50 | **P<0.0001** |
| **Figure 6c** | RRS x Treatment | F (1, 19) = 0.002195 | P=0.9631 |
| | Treatment | F (1, 19) = 2.825 | P=0.1092 |
| | RRS | F (1, 19) = 32.07 | **P<0.0001** |
| **Figure 6d** | Time | F (5, 95) = 321.6 | **P<0.0001** |
| | Treatment | F (1, 19) = 0.8504 | P=0.3680 |
| | RRS | F (1, 19) = 18.18 | **P=0.0004** |
| | Time x Treatment | F (5, 95) = 0.2603 | P=0.9336 |
| | Time x RRS | F (5, 95) = 6.212 | **P<0.0001** |
| | Treatment x RRS | F (1, 19) = 1.284 | P=0.2712 |
| | Time x Treatment x RRS | F (5, 95) = 0.3140 | P=0.9035 |
| **Figure 6e** | Time | F (5, 95) = 329.7 | **P<0.0001** |
| | Treatment | F (1, 19) = 0.3831 | P=0.5433 |
| | RRS | F (1, 19) = 11.58 | **P=0.0030** |
| | Time x Treatment | F (5, 95) = 0.3517 | P=0.8800 |

| | | | |
|---|---|---|---|
| | Time x RRS | F (5, 95) = 5.362 | **P=0.0002** |
| | Treatment x RRS | F (1, 19) = 0.08282 | P=0.7766 |
| | Time x Treatment x RRS | F (5, 95) = 0.2376 | P=0.9450 |
| **Figure 7a** | RRS x Treatment | F (1, 17) = 2.678 | P=0.1201 |
| | Treatment | F (1, 17) = 6.193 | **P=0.0235** |
| | RRS | F (1, 17) = 9.017 | **P=0.0080** |
| **Figure 7b** | RRS x Treatment | F (1, 21) = 6.731 | **P=0.0169** |
| | Treatment | F (1, 21) = 0.04469 | P=0.8346 |
| | RRS | F (1, 21) = 2.986 | P=0.0987 |
| **Figure 7c** | RRS x Treatment | F (1, 21) = 6.567 | **P=0.0181** |
| | Treatment | F (1, 21) = 7.871 | **P=0.0106** |
| | RRS | F (1, 21) = 1.829 | P=0.1906 |
| **Figure 7d** | Time | F (5, 105) = 372.6 | **P<0.0001** |
| | Treatment | F (1, 21) = 27.82 | **P<0.0001** |
| | RRS | F (1, 21) = 7.768 | **P=0.0110** |
| | Time x Treatment | F (5, 105) = 4.181 | **P=0.0017** |
| | Time x RRS | F (5, 105) = 2.086 | P=0.0729 |
| | Treatment x RRS | F (1, 21) = 15.86 | **P=0.0007** |
| | Time x Treatment x RRS | F (5, 105) = 1.613 | P=0.1629 |
| **Figure 7e** | Time | F (5, 105) = 412.5 | **P<0.0001** |
| | Treatment | F (1, 21) = 11.11 | **P=0.0032** |
| | RRS | F (1, 21) = 3.321 | P=0.0827 |
| | Time x Treatment | F (5, 105) = 3.723 | **P=0.0038** |
| | Time x RRS | F (5, 105) = 1.773 | P=0.1247 |
| | Treatment x RRS | F (1, 21) = 4.124 | P=0.0551 |
| | Time x Treatment x RRS | F (5, 105) = 1.653 | P=0.1524 |

**Supplementary Table 2: RNA Integrity Number (RIN) of samples used for RNAseq.**

| Sample name | RIN |
|---|---|
| SC1 | 8.4 |
| SC2 | 6.9 |
| SC4 | 5.8 |
| SC5 | 7.5 |
| SC6 | 8.6 |
| SC7 | 9.3 |
| SC8 | 7.5 |
| SC9 | 6.6 |
| SC10 | 9.5 |
| SC11 | 9 |
| SC12 | 7.3 |
| SC15 | 8.8 |
| SC17 | 9.2 |
| SC20 | 7.7 |
| SC21 | 9 |
| SC24 | 9 |
